# Selective protein O-GlcNAcylation in cells by a proximity-directed O-GlcNAc transferase

**DOI:** 10.1101/828921

**Authors:** Daniel H. Ramirez, Chanat Aonbangkhen, Hung-Yi Wu, Jeffrey A. Naftaly, Stephanie Tang, Timothy R. O’Meara, Christina M. Woo

## Abstract

O-Linked *N*-acetylglucosamine (O-GlcNAc) is a monosaccharide that plays an essential role in cellular signaling throughout the nucleocytoplasmic proteome of eukaryotic cells. Yet, the study of post-translational modifications like O-GlcNAc has been limited by the lack of strategies to induce O-GlcNAcylation on a target protein in cells. Here, we report a generalizable genetic strategy to induce O-GlcNAc to specific target proteins in cells using a nanobody as a proximity-directing agent fused to O-GlcNAc transferase (OGT). Fusion of a nanobody that recognizes GFP (nGFP) or a nanobody that recognizes the four-amino acid sequence EPEA (nEPEA) to OGT(4), a truncated form of OGT, yielded a nanobody-OGT(4) construct that selectively delivered O-GlcNAc to the target protein (e.g., JunB, cJun, Nup62) and reduced alteration of global O-GlcNAc levels in the cell. Quantitative chemical proteomics confirmed the selective increase in O-GlcNAc to the target protein by nanobody-OGT(4). Glycoproteomics revealed that nanobody-OGT(4) or full-length OGT produced a similar glycosite profile on the target protein. Finally, we demonstrate the ability to selectively target endogenous α-synuclein for glycosylation in HEK293T cells. Thus, the use of nanobodies to redirect OGT substrate selection is a versatile strategy to induce glycosylation of desired target proteins in cells that will facilitate discovery of O-GlcNAc functions and provide a mechanism to engineer O-GlcNAc signaling. The proximity-directed OGT approach for protein-selective O-GlcNAcylation is readily translated to additional protein targets and nanobodies that may constitute a generalizable strategy to control post-translational modifications in cells.

**Significance Statement:** Nature uses post-translational modifications (PTMs) like glycosylation as a mechanism to alter protein signaling and function. However, the study of these modified proteins in cells is confined to loss-of-function strategies, such as mutagenic elimination of the modification site. Here, we report a generalizable strategy for induction of O-GlcNAc to a protein target in cells. The O-GlcNAc modification is installed by O-GlcNAc transferase (OGT) to thousands of nucleocytoplasmic proteins. Fusion of a nanobody to OGT enables the selective increase of O-GlcNAc levels on a series of target proteins. The described approach will facilitate direct studies of O-GlcNAc and its regulatory enzymes and drive new approaches to engineer protein signaling via a strategy that may be conceptually translatable to additional PTMs.

## Introduction

All proteins are modified by post-translational modifications (PTMs) that control the protein lifetime, protein–protein interactions, and downstream signaling. However, methods to manipulate these modifications for evaluation of their functional output or engineering of cellular signals are limited. For example, 15% of the cellular proteome is modified by O-linked ß-*N*-acetyl glucosamine (O-GlcNAc), a PTM that consists of a single monosaccharide attached to serine or threonine residues on nuclear, cytosolic, and mitochondrial proteins. Due to the ubiquitous nature of the modification, O-GlcNAc has been implicated in numerous biological processes and diseases, including the immune response (1, 2), cancer progression (3), neurodegeneration (4), and diabetes (5, 6). However, the O-GlcNAc modification, like many cellular PTMs, cannot be tuned on particular glycoproteins within the cell. Global alteration of O-GlcNAc levels can be achieved through manipulating gene expression or chemical inhibitors (7, 8), but relating biological effects to a specific glycoprotein requires extensive follow-up studies. Elimination of the O-GlcNAc modification on specific target proteins is possible following glycosite mapping and mutagenic removal of the glycosite from the protein in cells. These methods have yielded defined functions for O-GlcNAc (3), but prevent analysis of competing PTM pathways (e.g., phosphorylation (9), ubiquitinylation(10)) and must be laboriously developed for every target protein if the exact glycosite is known. They are also challenging to implement for larger proteins or those carrying multiple glycosites. By contrast, a general method to selectively induce glycosylation on a desired target protein would enable the rapid and systematic evaluation of protein O-GlcNAcylation in cells.

In contrast to most PTMs, O-GlcNAc is installed and removed by only two enzymes: O-GlcNAc transferase (OGT) and O-GlcNAcase (OGA), which modify over 3,000 protein substrates (Figure 1) (11-15). O-GlcNAc is critical to cellular function as deletion of OGT in mice is embryonic lethal (16), deletion of OGA leads to perinatal death (17), and the conditional deletion of OGT in numerous cell types leads to senescence and apoptosis (18). OGT is a modular protein found in three major isoforms (ncOGT, mOGT, sOGT) consisting of a catalytic domain connected to a variable length tetratricopeptide repeat (TPR) domain (Figure 1B). The TPR domain is thought to primarily direct substrate and glycosite selection (19, 20), although the parameters that dictate how OGT and OGA dynamically regulate thousands of O-GlcNAc modification sites on substrates are still under investigation (21, 22). Given the dynamic nature of O-GlcNAcylation and the large number of substrates modified by OGT, we hypothesized that controlling O-GlcNAc levels in a protein-selective manner could be achieved through induced proximity (Figure 1C). Of the several mechanisms to induce protein–protein interactions (23), the defined properties of nanobodies were particularly attractive. In contrast to antibodies (∼150 kDa), nanobodies are small (∼12 kDa), highly-selective binding agents that are frequently used in affinity-based assays, imaging, X-ray crystallography, and recently as directing groups to recruit GFP fusion proteins for degradation or to a desired genomic loci (24-27). We thus sought to evaluate the potential for nanobodies to re-direct the activity of OGT and induce O-GlcNAcylation on a series of target proteins.

**Figure 1.**
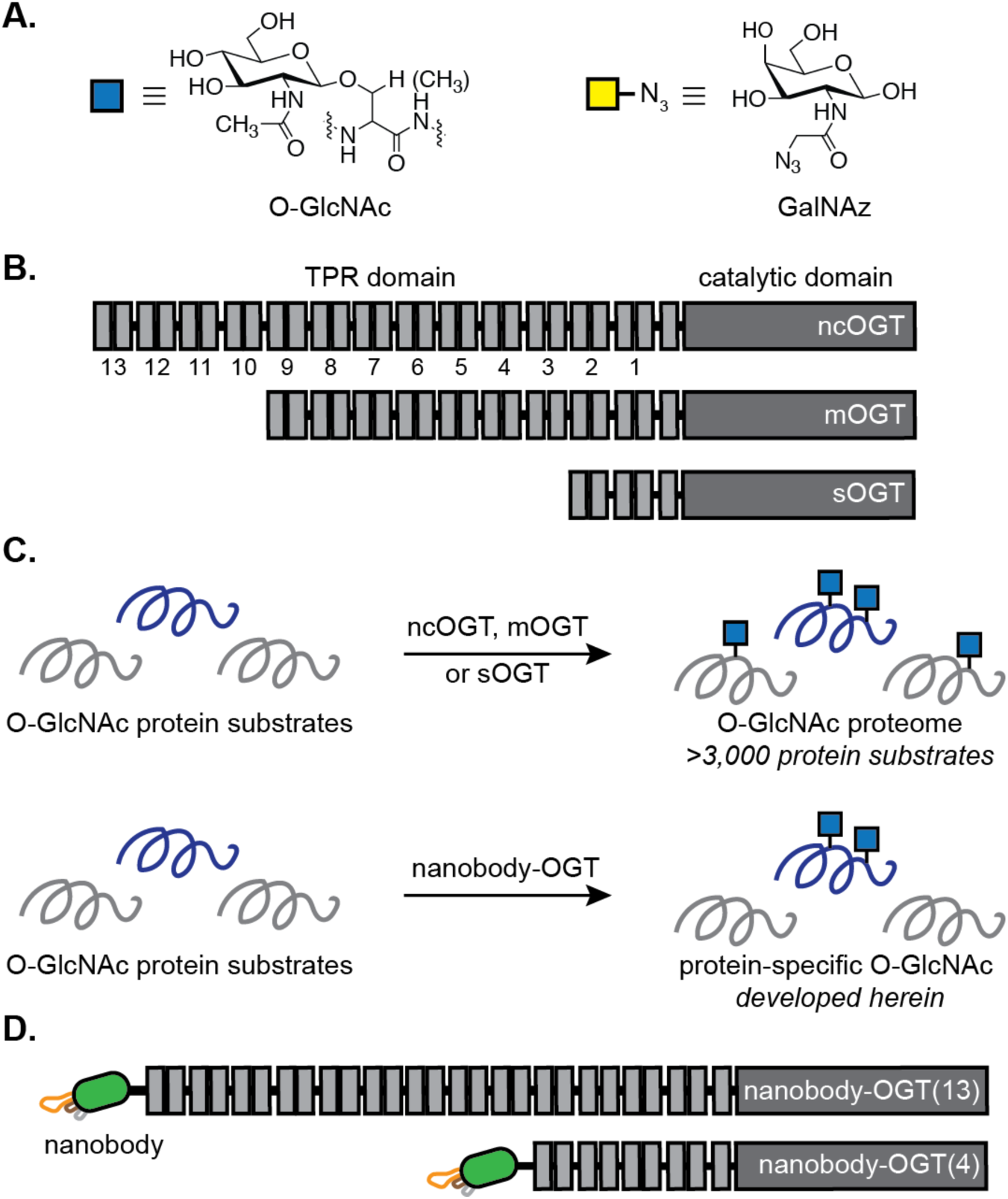
Schematic overview of proximity-directed OGT strategy. **A.** Structures of O-GlcNAc and GalNAz. **B.** Linear representation of natural OGT isoforms ncOGT, mOGT, and sOGT. **C.** Strategy for selective induction of O-GlcNAc using a proximity-directed nanobody-OGT to transfer O-GlcNAc to the target protein. **D.** Linear representation of nanobody-OGT(13) and nanobody-OGT(4) fusion proteins.

Here, we report the development of nanobody-OGT fusion proteins to yield the first proximity-directed OGT for elevation of O-GlcNAc levels on specific target proteins in cells (Figure 1D). We show that fusion of a nanobody to OGT induces O-GlcNAc to expressed or endogenous target proteins in the cell and that substitution of part of the TPR domain with the nanobody increased selectivity for the target protein. Protein-selective O-GlcNAcylation was achieved using a nanobody that recognizes GFP (nGFP) and a nanobody that recognizes EPEA, a four-amino acid sequence derived from α-synuclein (nEPEA). Fusion of these nanobodies to full-length OGT(13) and TPR-truncated OGT(4) induced O-GlcNAcylation to several co-expressed target proteins in HEK293T cells. Quantitative proteomics and glycoproteomics revealed selective induction of O-GlcNAc to the target protein by the nanobody-OGT fusion proteins with similar glycosite selectivity on evaluated target proteins JunB and Nup62, without broad perturbation of the global O-GlcNAc proteome. We finally demonstrate targeted glycosylation of endogenous α-synuclein in HEK293T cells by nEPEA-OGT(4). Thus, the use of nanobodies to proximity-direct OGT yields a versatile platform for protein-selective O-GlcNAcylation in live cells that will facilitate analysis of the essential functions of O-GlcNAc and enable new mechanisms to engineer the O-GlcNAc proteome.

## Results

### Design and expression of proximity-directed nanobody-OGT(13) constructs

Our overarching objective to engineer O-GlcNAc modifications in the cellular proteome inspired the design of initial nanobody-OGT fusion proteins using a GFP-recognizing nanobody, nGFP (28). We initiated our studies with nGFP fused to the full-length OGT that possesses 13 TPRs [residues 1–1046, nGFP(13)] and a RFP or GFP fusion to full-length OGT [RFP(13), GFP(13)] as an untargeted control for comparison to the nanobody-OGT construct due to similarity in expression levels (Figure 2A, Figure S1). All fusions to OGT were connected by an alpha-helical rigid linker (EAAAK)_4_. Expression of OGT(13) and RFP(13) was distributed throughout the cytoplasm and nucleus as shown by confocal fluorescence microscopy (Figure 2B). The nGFP(13) construct was distributed throughout the nucleocytoplasmic space in an analogous manner.

**Figure 2.**
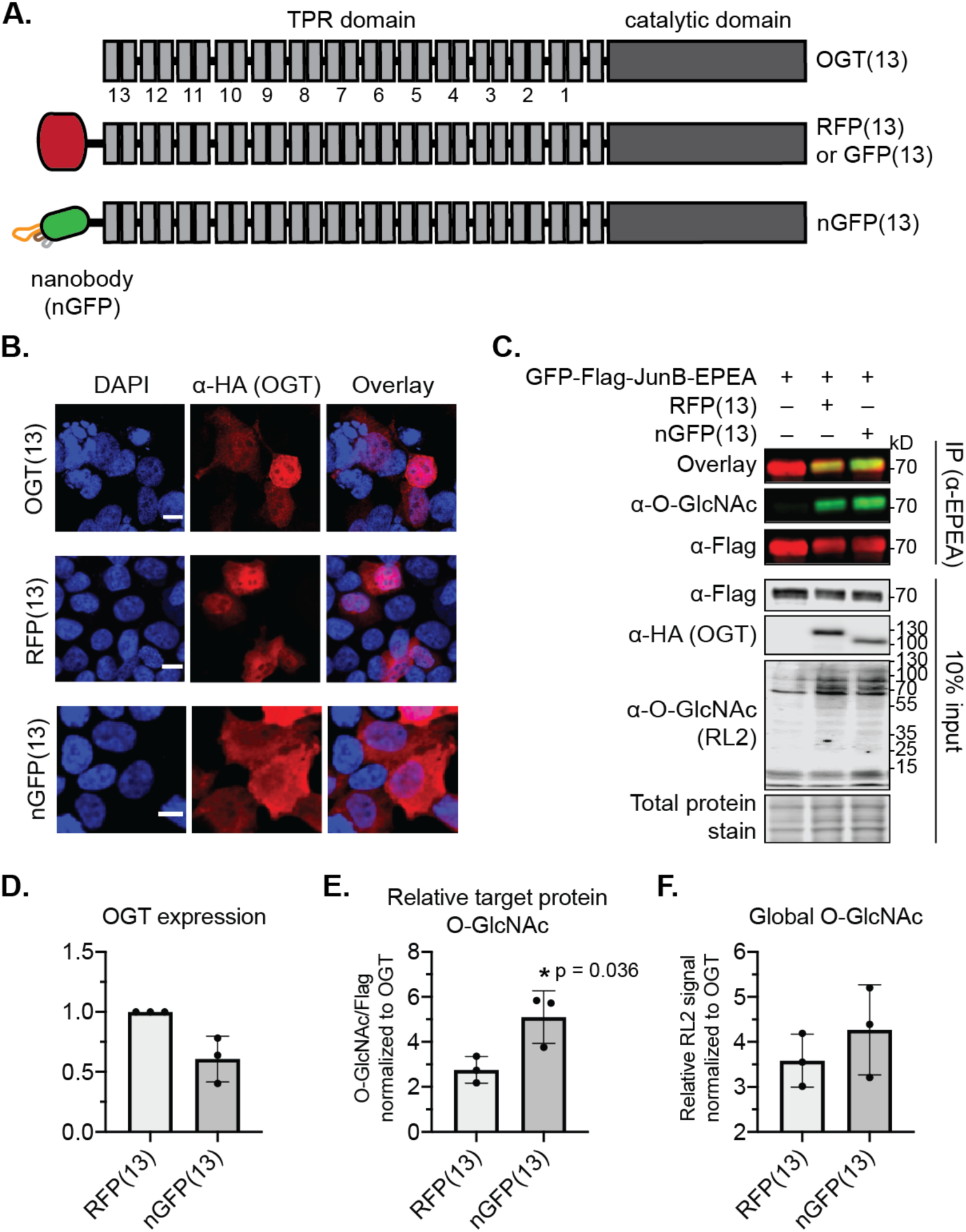
Characterization of nanobody-OGT(13) constructs as proximity-directing glycosyltransferase constructs. **A.** Linear representation of full-length OGT(13), RFP(13), and nGFP(13). **B.** Subcellular localization of OGT(13), RFP(13), and nGFP(13) constructs expressed in HEK293T cells by confocal fluorescent microscopy. Scale bars represent 20 µm. **C.** Western blot of O-GlcNAc levels on GFP-Flag-JunB-EPEA after immunoprecipitation with EPEA-beads from HEK293T cells. The expression of the various constructs was verified by Western blot analysis (10% input). **D.** Quantification of OGT expression. **E.** Quantification of O-GlcNAc levels on GFP-Flag-JunB-EPEA after normalization to OGT expression. **F.** Quantification of O-GlcNAc levels in whole cell lysate after normalization to OGT expression. Data are representative of three biological replicates per experiment. Error bars represent standard deviation, * represents a p-value <0.05 under a two-tailed t-test. Full blot and confocal images can be found in Figure S2, S3.

We next evaluated the proximity-directing ability of nGFP(13) in HEK293T cells co-expressing with GFP-Flag-JunB-EPEA, a transcription factor carrying multiple O-GlcNAc sites (15), as the target protein. Immunoprecipitation of GFP-Flag-JunB-EPEA and probing for O-GlcNAc revealed an increase in O-GlcNAc levels on the target protein that was dependent on the co-transfected nGFP(13) (Figure 2C). The O-GlcNAcylated target protein was significantly increased when co-expressed with nGFP(13) as compared to RFP(13) (Figure 2D, E). However, global O-GlcNAc levels were similarly elevated in the presence of RFP(13) and nGFP(13), implying that although nGFP(13) elevated levels of the O-GlcNAcylated target protein JunB, the selectivity for the target protein could be further improved (Figure 2F).

### Design and expression of proximity-directed nanobody-OGT(4) constructs

While pleased with the ability to proximity-direct nGFP-OGT(13) to a target protein in live cells, we hypothesized that reduction of the TPR domain of OGT would further improve the selectivity for the target protein and reduce global elevation of O-GlcNAc levels. We thus sought to evaluate a truncated OGT to optimize proximity-direction of OGT via the nanobody and used a truncated OGT with 4 TPRs due to prior structural characterization of the substrate binding mechanism (20). In addition, a unique advantage of using nanobodies to redirect enzymatic function is the ability to modularly fuse nanobodies that recognize specific target proteins within the cell. We thus evaluated OGT(4) fusions of nGFP and an additional nanobody, nEPEA (Figure 3A). The nEPEA nanobody was originally developed against α-synuclein and recognizes the four-amino acid EPEA tag as a fusion to the C-terminus of proteins (29). The EPEA tag was particularly appealing as the sequence itself cannot be glycosylated and is minimally perturbative to protein structure (29). However, to remove potential competition for the target protein by endogenous α-synuclein in HEK293T cells, we generated an α-syn KO HEK293T cell line for studies employing the nEPEA nanobody (Figure S4).

**Figure 3.**
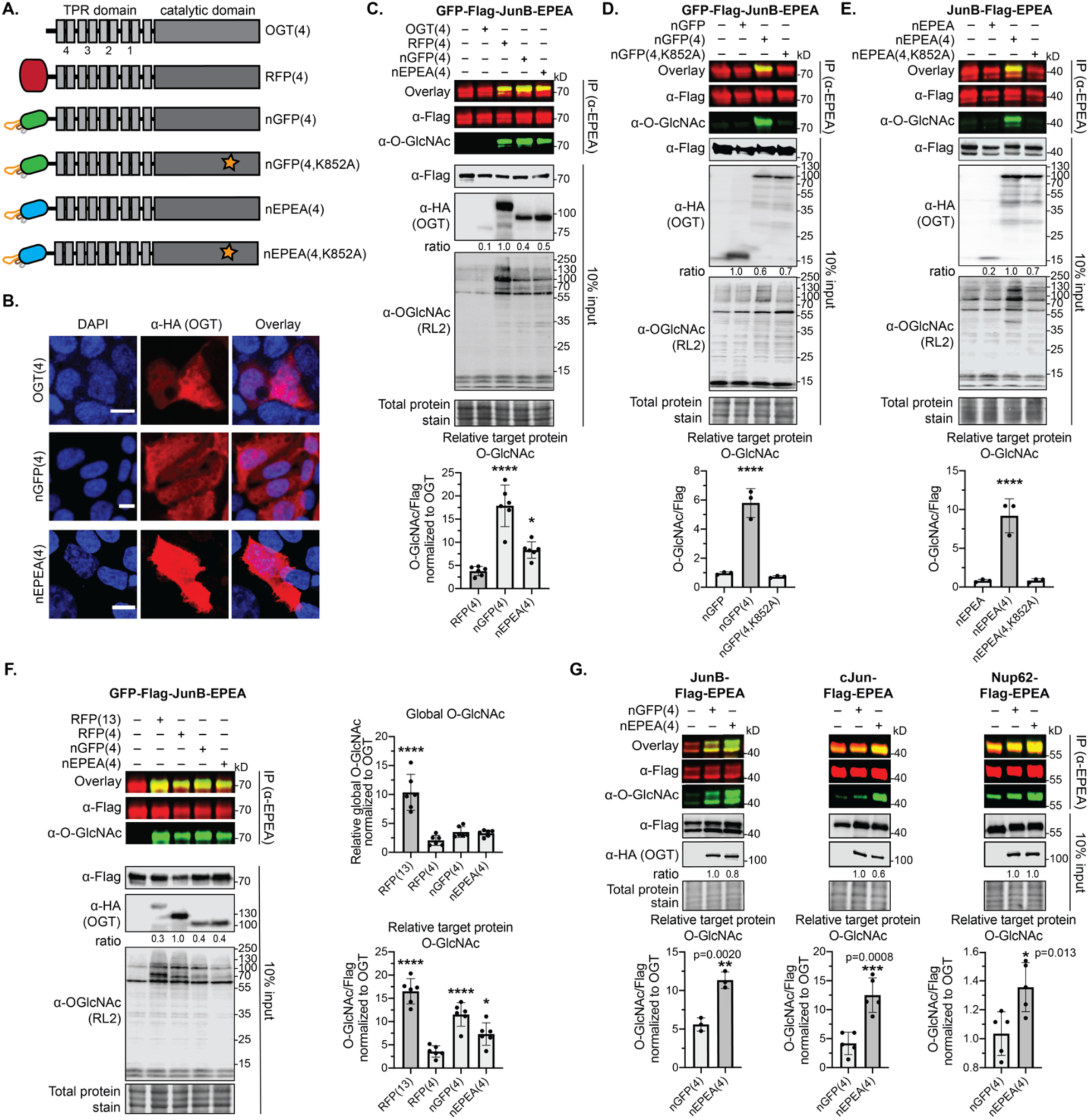
Characterization of nanobody-OGT(4) constructs as proximity-directing glycosyltransferase constructs. **A.** Linear representation of TPR truncated OGT(4), RFP(4), nGFP(4), nEPEA(4), and catalytically inactive mutants. **B.** Subcellular localization of OGT(4), nGFP(4), and nEPEA(4) in HEK293T cells by confocal fluorescent microscopy. Scale bars represent 20 µm. **C.** Western blot and quantification of O-GlcNAc levels on GFP-Flag-JunB-EPEA after immunoprecipitation with EPEA-beads. The expression of the various constructs was verified by Western blot analysis (10% input). **D.** Western blot and quantification of O-GlcNAc levels on GFP-Flag-JunB-EPEA after immunoprecipitation with EPEA-beads. The expression of the various constructs was verified by Western blot analysis (10% input). **E.** Western blot and quantification of O-GlcNAc levels on JunB-Flag-EPEA after immunoprecipitation with EPEA-beads. The expression of the various constructs was verified by Western blot analysis (10% input). **F.** Western blot and quantification of O-GlcNAc levels on GFP-Flag-JunB-EPEA after immunoprecipitation with EPEA-beads. The expression of the various constructs was verified by Western blot analysis (10% input). **G.** Western blot and quantification of O-GlcNAc levels on JunB-Flag-EPEA, cJun-Flag-EPEA, and Nup62-Flag-EPEA after immunoprecipitation with EPEA-beads from α-syn KO HEK293T cells co-transfected with the indicated nanobody-OGT fusion protein and target protein. The expression of the various constructs was verified by Western blot analysis (10% input). At least three biological replicates were performed per experiment. Error bars represent standard deviation, * represents p ≤ 0.05, ** represents p ≤ 0.01, *** represents p ≤ 0.001, **** represents p ≤ 0.0001 under a two-tailed t-test or one-way ANOVA. Full blot and confocal images can be found in Figures S5–S10.

Expression of the OGT(4) fusion proteins in α-syn KO HEK293T cells showed a subcellular localization throughout the nucleocytoplasmic space by confocal fluorescence microscopy (Figure 3B). The activity of OGT(4), RFP(4), nGFP(4) and nEPEA(4) was evaluated against the same target protein GFP-Flag-JunB-EPEA (Figure 3C). Both nGFP(4) and nEPEA(4) significantly increased the O-GlcNAcylated target protein relative to the untargeted controls OGT(4) and RFP(4). To determine if the activity observed came from our nanobody-OGT(4) fusion, we generated two additional controls consisting of either the nanobody (nGFP, nEPEA) or a catalytically-inactive version of the nanobody-OGT(4) that is unable to bind to UDP-GlcNAc [nGFP(4,K852), nEPEA(4,K852A)] (30). Increased levels of O-GlcNAcylated GFP-Flag-JunB-EPEA are installed directly from the active nGFP(4), but not the nanobody nGFP alone or a catalytically inactive mutant nGFP(4,K852A) (Figure 3D). Similarly, O-GlcNAcylated JunB-Flag-EPEA is specifically increased on co-expression of nEPEA(4), but not in the presence of nEPEA or the catalytically inactive nEPEA(4,K852A) (Figure 3E). The O-GlcNAcylation activity of the OGT(4) fusions was further compared to the full-length RFP(13) (Figure 3F). As expected, truncation of the TPR domain as in OGT(4) attenuated the increase in global O-GlcNAc levels observed with RFP(13). O-GlcNAc activity for the desired target protein could therefore be selectively reinstated by use of the nanobody for proximity direction (Figure 3F).

The versatility of the nanobody-OGT(4) system to selectively increase the O-GlcNAcylated target protein was further evaluated against three targets: JunB-Flag-EPEA, cJun-Flag-EPEA, and Nup62-Flag-EPEA in HEK293T cells. We found that the O-GlcNAcylated target protein significantly increased under proximity-direction of the matched nEPEA(4), but not the mismatched nGFP(4) in each case (Figure 3G). Collectively, these data point to the successful increase in selective O-GlcNAcylation by replacing elements of the TPR domain with the nanobody and the modular ability of the nanobody-OGT(4) to increase O-GlcNAc levels using GFP- or EPEA-tagged target proteins.

### Quantitative proteomics with expressed nanobody-OGT constructs

To better quantify the selectivity of the nanobody-OGT constructs for the target protein, we performed a series of quantitative proteomics experiments by mass spectrometry (MS). Cellular lysates were collected following co-expression in α-syn KO HEK293T cells of nEPEA(4) with JunB-Flag-EPEA as the target protein. Then, lysates were chemoenzymatically labeled with UDP-GalNAz to introduce an azido-sugar for copper-catalyzed azide-alkyne cycloaddition (CuAAC) with a biotin-azide probe and enrichment on streptavidin–agarose beads (31). The O-GlcNAcylated proteins were digested on-bead and labeled with Tandem Mass Tags (TMT) for MS analysis. Glycoprotein enrichment was determined relative to the control for high-confidence proteins [number of unique peptides ≥ 2, 1% false discovery rate (FDR)] (Figure 4A, Table S1). These data show that JunB-Flag-EPEA was the only protein enriched >4-fold in samples co-expressing nEPEA(4), implying good target selectivity by the nEPEA nanobody (highlighted in red and blue, respectively, Figure 4A).

**Figure 4.**
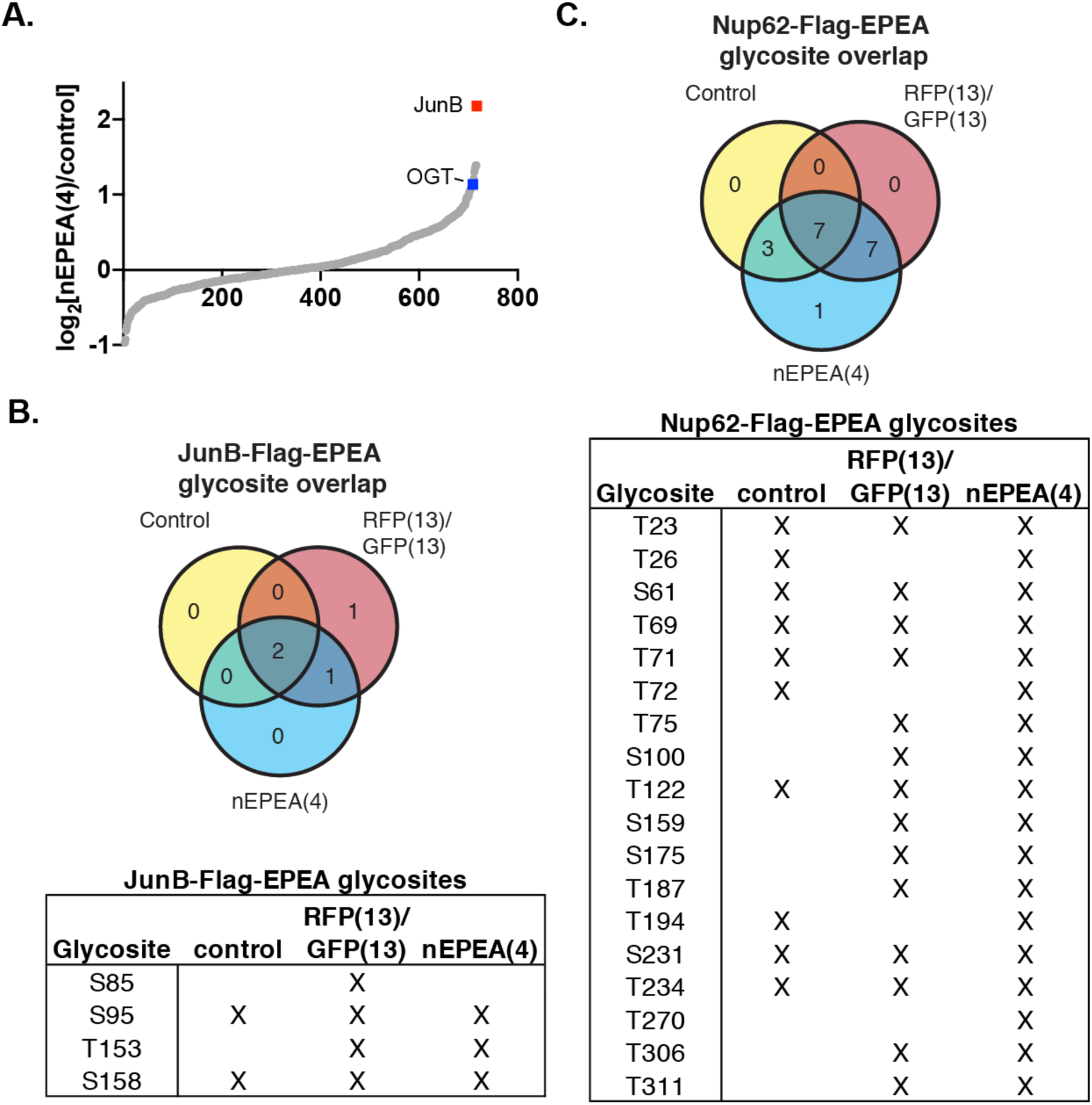
Quantitative proteomics and glycoproteomics of the global O-GlcNAc proteome from α-syn KO HEK293T cells. **A.** Quantitative proteomics of enriched O-GlcNAcylated proteins from α-syn KO HEK293T cells after co-expression of nEPEA(4) (highlighted in blue) and JunB-Flag-EPEA (highlighted in red). **B.** Venn diagram and list of unambiguous glycosite assignments for JunB-Flag-EPEA. **C.** Venn diagram and list of unambiguous glycosite assignments for Nup62-Flag-EPEA. The target protein was co-expressed with the indicated OGT fusion protein in α-syn KO HEK293T cells, immunoprecipitated, and analyzed by MS. Only mono-glycosylated peptides with unambiguous assignments and a PSM count > 2 were given an unambiguous glycosite designation. Three biological replicates were performed per experiment.

### Characterization of glycosites produced by proximity-directed OGT constructs

We next sought to address whether the TPR-truncated OGT(4) exhibited glycosite selectivity that was broadly similar to ncOGT or displayed activity that may be more akin to sOGT against protein targets in cells. Previous in vitro structural work on the mechanism of OGT substrate binding implicated a series of asparagine and aspartate residues lining the first four TPRs of OGT to be responsible for binding and modifying a large percentage of the O-GlcNAc proteome (32, 33). Since these residues are conserved in OGT(4), we sought to characterize the protein regions and the associated glycosites installed by the nanobody-OGT(4) as compared to the full-length OGT(13). JunB-Flag-EPEA was expressed alone as the control or co-expressed with an undirected full-length OGT [RFP(13) or GFP(13)] or nEPEA(4) in α-syn KO HEK293T cells and was affinity purified, digested with chymotrypsin, and analyzed by MS in biological triplicate. Unambiguous glycosites were determined based on previously established criteria (15). Four unambiguous glycosites were identified from JunB-Flag-EPEA. Three of the four unambiguous glycosites (S95, T153, S158) were identified in at least one nEPEA(4) sample and one reference sample [control, RFP(13)/GFP(13)] (Figure 4B, Table S2). The remaining glycosite S85 was identified only in samples co-expressing GFP(13). This data indicate that replacement of parts of the TPR domain with the nanobody results in a fusion construct that targets broadly similar glycosites as a full-length OGT(13).

We further evaluated the glycosite specificity of nEPEA(4) on the highly O-GlcNAcylated protein Nup62. A total of 18 unambiguous glycosites were mapped to Nup62-Flag-EPEA (Figure 4C, Table S3). Of these 18 glycosites, 17 glycosites were found in at least one of the reference samples and in one of the samples expressing nEPEA(4). The remaining glycosite (T270) was observed only on co-expression of nEPEA(4). Seven glycosites (T75, S100, S159, S175, T187, T306, T311) on Nup62-Flag-EPEA were found only on co-expression of an OGT construct. Taken together, nEPEA(4) displayed analogous glycosite selectivity towards Nup62-Flag-EPEA and JunB-Flag-EPEA while increasing overall levels of the O-GlcNAcylated target protein.

### Targeting endogenous α-synuclein for O-GlcNAcylation with proximity-directed OGT

A major advantage of using nanobodies for OGT recruitment is the potential to recruit OGT to endogenous target proteins. Since the nEPEA nanobody was originally developed against the C-terminal EPEA sequence of α-synuclein, we evaluated whether nEPEA(4) could increase glycosylation of endogenous α-synuclein in a selective manner. Endogenous α-synuclein glycosylation was visualized using a mass-shift assay, which uses a chemical reporter for O-GlcNAc to install a PEG5K tag for determination of O-GlcNAc protein stoichiometry (Figure 5A) (34). As expected, we observed an increase in O-GlcNAc levels on α-synuclein by mass shift assay on expression of RFP(13) and nEPEA(4), but not the catalytically inactive mutant nEPEA(4,K852A) (Figure 5B). Notably, expression of the untargeted RFP(13) resulted in a dramatic increase in O-GlcNAcylation across the global HEK293T cell proteome that was significantly reduced in the nEPEA(4) sample. Comparison of the glycosylated α-synuclein to the global levels of O-GlcNAc allowed us to obtain a selectivity factor for the target protein over the global O-GlcNAc proteome by nEPEA(4). Based on the selectivity factor, nEPEA(4) enables a significant increase in O-GlcNAcylated α-synuclein with reduced perturbations to the global O-GlcNAc levels observed as compared to expression of an untargeted OGT control, such as RFP(13). Taken together, these data demonstrate the high selectivity and versatility of proximity-directed nanobody-OGT(4) constructs to transfer O-GlcNAc to the desired target protein, either as a tagged or endogenous target protein, with reduced impact on global O-GlcNAc levels.

**Figure 5.**
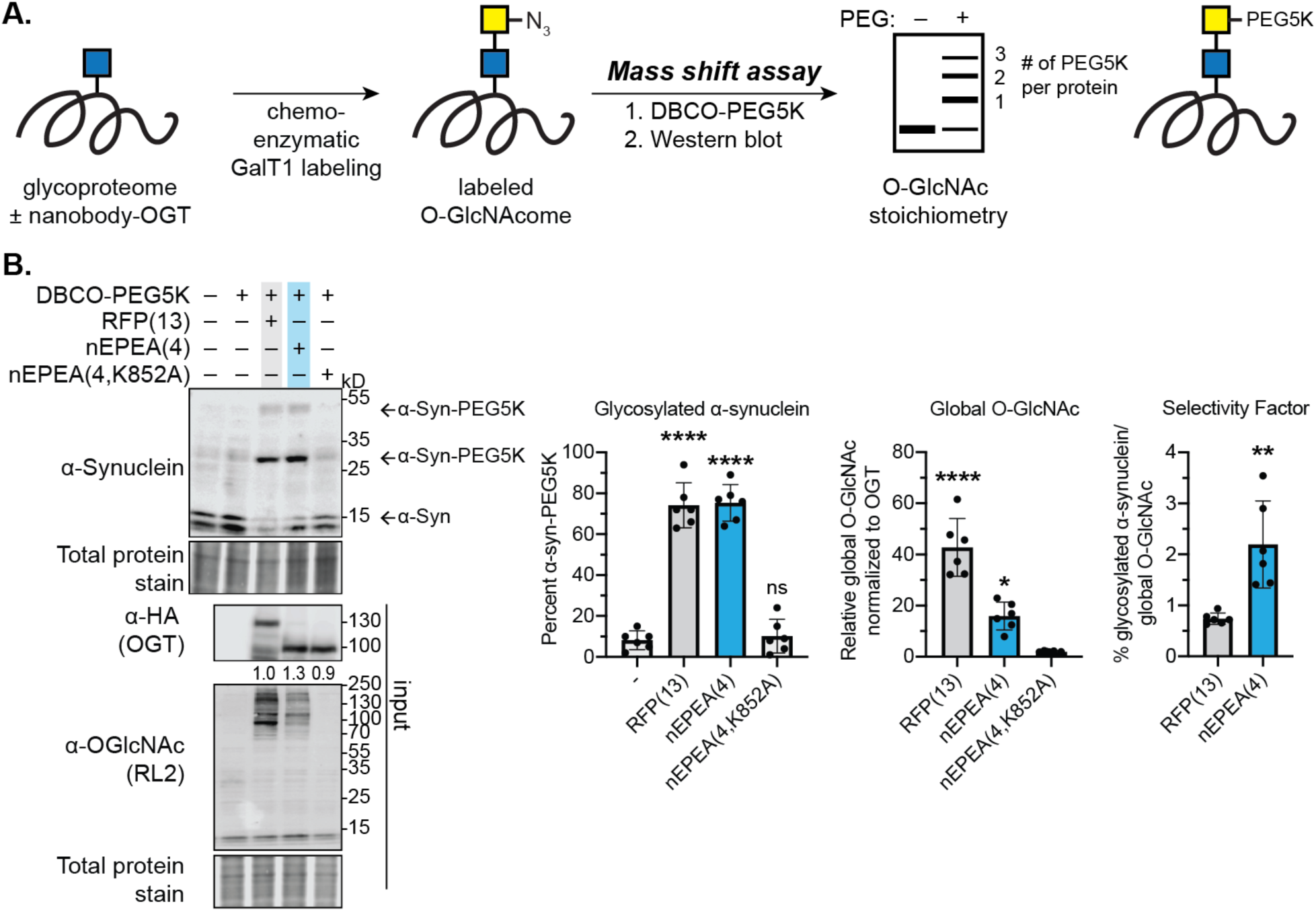
Proximity-directed nanobody-OGT constructs induce O-GlcNAc on endogenous α-synuclein target proteins. **A.** Mass-shift assay workflow. O-GlcNAcylated cell lysates are chemoenzymatically labeled with GalNAz using GalT1. The GalNAz is reacted with a DBCO-PEG5K and a western blot is performed to obtain an O-GlcNAc stoichiometry. **A.** Western blot and quantification of O-GlcNAc induced to α-synuclein by a mass shift assay. The indicated nanobody-OGT construct was expressed in HEK293T cells, the cells were lysed, chemoenzymatically labeled, and analyzed by mass shift assay. Global O-GlcNAc levels and the expression of the nanobody-OGT constructs was verified by Western blot analysis (10% input). At least six biological replicates were performed per experiment. Error bars represent standard deviation, ns represents p ≥ 0.05, * represents p ≤ 0.05, ** represents p ≤ 0.01, **** represents p ≤ 0.0001 under a two-tailed t-test or one-way ANOVA. Full blots can be found in Figure S11

## Discussion

We report here the ability to increase O-GlcNAcylation on a target protein in live cells through fusion of a nanobody to OGT. The study of PTMs like O-GlcNAc in cells typically rely on global alteration of the modification or loss-of-function genetic mutagenesis strategies to eliminate the modification site. Alternatively, a gain-of-function approach to selectively induce the modification in cells would provide an intriguing opportunity to evaluate the effect of the modification on the protein and introduce a mechanism to selectively engineer cellular signaling pathways. Proximity-directed OGT constructs were designed by fusion of nGFP or nEPEA to full-length OGT(13) or TPR-truncated OGT(4) connected by a rigid linker (EAAAK)_4_. These constructs were successfully recruited to target proteins carrying either GFP or the four-amino acid sequence EPEA for transfer of O-GlcNAc. We demonstrate that all three nanobody-OGT constructs directly increase O-GlcNAcylation to target proteins and show that replacement of part of the TPR domain of OGT with a nanobody can significantly improve selectivity for the target protein as compared to an untargeted control (Figure 2, 3). Although truncation of the TPR domain has been reported to reduce glycosyltransferase activity for proteins, presumably through the loss of binding to the protein substrate (21, 35), the activity for the target protein is selectively regained through the nanobody’s affinity for the target protein. The nEPEA-OGT(4) fusion protein was further validated by quantitative proteomics and glycoproteomics to establish the selective O-GlcNAcylation of the target protein without broad perturbation of the O-GlcNAc proteome at glycosites that are largely analogous to those installed by OGT(13) (Figure 4). The replacement of protein-recognition domains by a nanobody, established here with OGT, has broader implications for the design of additional proximity-directed enzymes for targeted installation of a range of protein modifications in live cells and illuminates a potential strategy to evaluate the glycosite selectivity of the TPR domain of OGT in cells. Emerging insight to how the TPR domain dictates substrate and glycosite selectivity highlights the need for further study in cells and their implications on the OGT isoforms mOGT and sOGT (32, 33).

Finally, we demonstrate the selective glycosylation of endogenous α-synuclein by nEPEA(4) (Figure 5). As increased glycosylation of α-synuclein has a protective effect on aggregation (36), the ability to induce O-GlcNAc on α-synuclein may have further implications in the treatment of Parkinson’s disease. Although nanobodies have been used for detection of endogenous target proteins (26), to our knowledge, the use of nanobody fusion proteins to recruit enzymatic activity to an endogenous target protein is first demonstrated here. The selective delivery of O-GlcNAc directly to endogenous target proteins will continue to expand as nanobodies that recognize additional native nucleocytoplasmic proteins are developed. We envision that the strategic incorporation of nanobodies that recognize specific protein conformations, or post-translational modification states, may be further employed to distinguish between co-translational or post-translational glycosylation and O-GlcNAc dynamics. For example, the intriguing shifts in the glycosylation pattern of Nup62 on elevated O-GlcNAcylation (S100–T311) may reflect broader regulatory properties of O-GlcNAc on domains of Nup62 (Figure 4C) (10). The design of additional nanobody-OGT constructs, with a broader range of TPR domain lengths, and evaluation of the resulting glycosite selectivity may lead to constructs that are protein-selective with enhanced glycosite or sub-sequence selectivity.

The use of nanobodies for proximity-directing groups has major advantages, including the relatively rapid evaluation of O-GlcNAcylation of new target proteins and the ability to target endogenous proteins, as compared to other induced proximity systems (e.g., chemically induced dimerization) (23, 37, 38). However, current nanobody systems must be controlled at the genetic level and cannot be chemically controlled after expression, though chemical stabilization of protein levels can be engineered (39). The use of nanobodies for induced proximity may reinforce artifactual interactions with the target protein that could disrupt normal protein interactions or cellular functions. To control for these potential effects, controls such as transfection with the nanobody or a catalytically inactive fusion of a nanobody [e.g., nEPEA(4,K852A)] should be included. In the future, optimization of binding parameters to tune reversibility may be possible using the several GFP-recognizing nanobodies that have been described (28), which may additionally provide insight to the properties of the TPR domain of OGT in cells. Co-expression of the nanobody-OGT and the target protein is also liable to spurious results simply due to artifacts from protein over-expression (40). Furthermore, as nanobody-OGT installs O-GlcNAc to the target protein in a catalytic manner, higher expression levels of the target protein is desirable. Future development of stable cell lines, inducible expression systems, or CRISPR knock-in strategies for introduction of peptide or protein tags may address these concerns.

In conclusion, we report a method for protein-selective O-GlcNAcylation using the first proximity-directed nanobody-OGT constructs. The approach enables the induction of O-GlcNAc to specific target proteins in cells for mechanistic studies of O-GlcNAcylated proteins. Analogously, substitution of protein domains that mediate substrate selection, like the TPR domain of OGT, with a nanobody is poised to enable the systematic measurement of the isolated functions of these domains in cells such as dissecting the scaffolding and catalytic functions of OGT. As a number of glycosyltransferases are composed of a lectin domain fused to a catalytic domain (41), nanobody fusion proteins possess a great potential for proximity-induced glycosylation of a broader array of glycan structures. The strategy reported here could drive new insights to the substrate selection mechanisms and functions of the glycoproteome and design principles for extension to the control of additional PTMs in a live cell environment.

## Acknowledgements

We thank Mr. Paul Schwein, Dr. Yun Ge, and Dr. Bo Yang for helpful discussions; and Dr. Bogdan Budnik, the director of the Harvard University Proteomics Facility, and Dr. Douglas Richardson, the director of Harvard Center of Biological Imaging (HCBI). Support from the National Institutes of Health (U01CA242098-01, C.M.W.), Burroughs Welcome Fund, Career Award at the Scientific Interface (C.M.W.), Milton Fund (C.M.W.), Mizutani Foundation for Glycoscience (C.M.W.), Sloan Foundation (C.M.W.), Merck Fellowship Fund, Corning Fund for Faculty Development, and Harvard University is gratefully acknowledged.

## Data Availability Statement

Mass spectrometry data is available via ProteomeXchange with identifier PXD016041.

Reviewer account details:

Username

Password: vdLqzQNF

## Online Methods

Commercial chemical materials, solvents, and reagents were used as received. Details about materials, reagents, antibodies, plasmids, geneblocks, and primers can be found in the Supplementary Information.

### Instrumentation

Protein quantification by bicinchoninic acid assay was measured on a multi-mode microplate reader FilterMax F3 (Molecular Devices LLC, Sunnyvale, CA). Cell lysis was performed using a Branson Ultrasonic Probe Sonicator (model 250). Fluorescence and chemiluminescence measurements were detected on an Azure Imager C600 (Azure Biosystems, Inc., Dublin, CA). All glycoproteomics data were obtained on a Waters ACQUITY UPLC connected in line to an Orbitrap Fusion Tribrid (ThermoFisher) within the Mass Spectrometry and Proteomics Resource Laboratory at Harvard University. Confocal fluorescence microscopy was performed at the Harvard Center for Biological Imaging (HCBI) using a Zeiss laser scanning confocal microscope (LSM) 880.

### Molecular cloning procedures

Plasmid #1 was a GFP nanobody fusion to full-length OGT developed by Gibson assembly and inserted into the pcDNA3.1 vector. The forward primer #1, used to amplify nGFP from cloning plasmid #1, contained an overlapping region to the pcDNA3.1 vector, Kozak sequence, a HA-tag for immunodetection, a Sgfl restriction enzyme (RE) site, and nucleotides complementary to the nGFP sequence. The reverse primer #2 contained complementary nucleotides to the nGFP sequence, a Sgsl RE site, and a stretch of nucleotides coding for a rigid helical linker composed of four iterations of the amino acid sequence EAAAK. The forward primer #3, used to amplify the OGT gene from cloning plasmid #2 included an overlapping region to the EAAAK linker, a BamH1 RE site, and complementary nucleotides to the OGT gene. The reverse primer #4 for OGT contained complementary nucleotides to the C-terminus of the OGT gene, a Not1 RE site, and overlapping nucleotides to the pcDNA3.1 vector. The pcDNA3.1 vector was restriction enzyme digested with HindIII and NotI enzymes and a Gibson Assembly was performed to construct the nGFP(13) plasmid #1.

Plasmids #2 and 3 were derived from plasmid #1 by restriction enzyme cloning by designing forward primers #5 and 6 containing a Sgfl RE site and complementary regions of interest in GFP or RFP and reverse primers #7 and 8 containing a Sgfl RE site and complementary regions to the C-terminus of GFP or RFP. PCR products were inserted into a Sgfl and Sgsl digested plasmid #1.

The OGT(13) plasmid #4 without the nanobody was created by designing a forward primer #9 containing a HindIII RE site, a HA tag, a BamHI RE site and complementary regions to OGT. The reverse primer #10 contained a NotI RE site and complementary regions to the C-terminus of OGT. PCR products were inserted into a HindIII and NotI digested pcDNA3.1 plasmid.

OGT(4) plasmids #5-8 were developed by restriction enzyme cloning by designing a forward primer #11 containing a BamHI RE site and complementary regions of interest in OGT and the reverse primer #10 containing a NotI RE site and complementary regions to the C-terminus of OGT. PCR products were inserted into BamHI and NotI digested plasmids #1, 2, 4 and 5.

OGT(K852A) plasmids #9 and 10 were developed by site-directed mutagenesis by designing forward primer #12 and reverse primer #13. Whole plasmid PCR products of plasmids #7 and 8 were obtained and blunt end cloning was performed.

GFP-Flag-JunB-EPEA plasmid #11 was developed by Gibson Assembly and inserted into the pcDNA3.1 vector. The forward primer #14, used to amplify GFP from cloning plasmid #3, contained an overlapping region to the pcDNA3.1 vector, a HindIII RE site, and nucleotides complementary to the GFP sequence. The reverse primer #15 contained complementary nucleotides to the GFP sequence, a Flag tag and one iteration of the amino acid sequence EAAAK. The forward primer #16, used to amplify the JunB gene from cloning plasmid #5 included an overlapping region to the Flag-EAAAK linker and complementary nucleotides to the JunB gene. The reverse primer #17 for JunB contained complementary nucleotides to the C-terminus of JunB, an EPEA tag, a XhoI RE site, and overlapping nucleotides to the pcDNA3.1 vector. The pcDNA3.1 vector was restriction enzyme digested with HindIII and XhoI enzymes and a Gibson Assembly was performed.

Nup62-Flag-EPEA plasmid #12 was developed by restriction enzyme cloning. A forward primer #18 containing a HindIII and Sgfl RE sites and a region complementary to the N-terminus of Nup62 was created. A reverse primer #19 with an XhoI RE site and regions complementary to the Flag and EPEA tag was created. The pcDNA3.1 vector was digested with the HindIII and XhoI restriction enzymes and restriction enzyme cloning was performed to develop the Nup62-Flag-EPEA plasmid #12. All other plasmids containing target proteins (plasmids #13 and 14) were created by designing forward and reverse primers containing either Sgfl or Sgsl RE sites and complementarity to the gene of interest and inserted into a Sgfl-and Sgsl-digested Nup62-Flag-EPEA plasmid #12.

EPEA nanobody gene blocks were amplified using primers #22 and 23, and EPEA nanobody constructs were created as with the nGFP fusions.

### Generation of α-synuclein CRISPR knockout HEK cell line

The α-synuclein CRISPR/Cas9 KO plasmid (human, Cat # sc-417273) and α-synuclein homology-directed DNA repair (HDR) plasmid (human, Cat # sc-417273-HDR) were purchased from Santa Cruz Biotechnology and transfected following the manufacturer’s instructions. The media was replaced with fresh DMEM growth media after 24 h. After 48 h of transfection, DMEM media supplemented with 2 µg/mL puromycin was added to the cells for KO-positive selection. The puromycin selection continued for 14 d with increasing concentration of puromycin up to 6 µg/mL prior to FACS to enrich for the RFP-positive cells (top 5% highest RFP intensity).

### Cell culture, transfection protocols, and cell lysate collection

Unless otherwise noted, the experiments were performed with HEK293T cells (a gift from the Bertozzi lab, Stanford University), or α-syn KO HEK293T cells. Cells were cultured in high-glucose with pyruvate Dulbecco’s Modified Eagle Medium (DMEM, Cat # 11995073) supplemented with 10% FBS and 1% penicillin–streptomycin at 37 °C in a humidified atmosphere with 5% CO_2_.

Samples for Western blot or immunofluorescence were prepared from cells seeded in a well of a sterile 6-well plate (VWR, Cat # 10062-892) at a density ∼1 × 10^6^ cells/well and transfected at ∼80% confluency the next day. For mass spectrometry-based glycoproteomics experiments, cells were seeded at the density ∼18 × 10^6^ cells/plate in sterile 150 mm tissue culture dishes (Corning, Cat # 25383-103) and transfected at ∼80% confluency the next day. Transient expression of the indicated proteins was performed by transfection with the desired plasmids following the manufacturer’s protocol. For immunofluorescence experiments, Lipofectamine 2000 (ThermoFisher, Cat # 11668027) was used at a ratio of 2 μg plasmid DNA to 5 µL of Lipofectamine. For all other experiments, TransiT-PRO (Mirus Bio, Cat # MIR 5740) was used with a ratio of 1 μg plasmid DNA to 1 µL of TransIT-PRO. As recommended by the manufacturers, transfection reagent and plasmid were diluted in Opti-MEM reduced serum medium (ThermoFisher, Cat # 31985070) during the transfection protocol. Cells were incubated for 36– 48 h after transfection before collection or visualization.

After 36–48 h of transfection, cells were collected and lysed by probe sonication in lysis buffer [150 µL of 2% SDS + 1× PBS + 50 µM Thiamet-G + 1× Protease inhibitors (cOmplete™, EDTA-free Protease Inhibitor Cocktail, Sigma Aldrich; Cat # 11873580001)]. A BCA assay was performed to determine protein concentration and the concentration was adjusted to 2.5 µg/µL with lysis buffer.

### EPEA-tag immunoprecipitation

For EPEA-tag immunoprecipitation and Western blot, α-syn KO HEK293T cells transfected in a 6-well plate were collected in lysis buffer [150 µL of 2% SDS + 1× PBS + 50 µM Thiamet-G + 1× Protease inhibitors (cOmplete™, EDTA-free Protease Inhibitor Cocktail, Sigma Aldrich; Cat # 11873580001)]. Samples were heated for 5 min at 95 °C and lysed by probe tip sonication 10 secs 10% amplitude. A BCA assay was performed to determine protein concentration and the concentration was adjusted to 2.5 µg/µL with lysis buffer. Protein lysate (100 µg) was incubated with C-tag resin (40 µL, Thermo Fisher; Cat # 191307005) and 1× PBS (500 µL). The mixture was incubated 12 h at 4 °C. The beads were washed 5× with 1× PBS (1 mL) and resuspended in 1× Laemmli sample buffer (50 µL; final concentrations: 60 mM Tris–HCl, 2% SDS, 10% glycerol, 5% ß-mercaptoethanol, 0.01% bromophenol blue) and heated for 5 min at 95 °C before loading on a gel for Western blot analysis.

For target protein glycoproteomics, α-syn KO HEK293T cells were plated in a 150-mm dish and were transfected with the corresponding plasmids for 48 h. Cells were collected in 2% SDS + 1× PBS + 1× Protease inhibitors + 50 µM Thiamet-G (1 mL) and heated for 5 min at 95 °C. Cells were lysed by probe tip sonication (30 sec, 15% amplitude). Samples were reduced with 25 mM DTT and heating for 5 min at 95 °C. Samples were then alkylated with 50 mM iodoacetamide for 1 h in the dark at 24 °C. Samples were precipitated by the addition of methanol (1.2 mL), chloroform (400 µL), and H_2_O (900 µL), vortexing, and centrifugation (10 min, 10,000 × g). Aqueous upper layer was discarded and methanol (1 mL) was added, sample was vortexed, and centrifuged (10 min, 10,000 × g). Sample was allowed to air dry (5 min) before resuspension in 2% SDS + 1× PBS (500 µL) by probe tip sonication. A BCA assay was performed and protein concentration was adjusted to 5 µg/µL with lysis buffer. Protein lysate (2.5 mg) was incubated with C-tag XL (300 µL, Thermo Fisher; Cat # 2943072005) and 1× PBS (1 mL). The mixture was incubated 12 h at 4 °C. The beads were washed 10× with 1× PBS (1 mL) then the beads were resuspended in 100 mM Tris-HCl + 10 mM CaCl_2_ (pH 8.0, 520 µL) for chymotrypsin digestion. Fifty µL of this mixture was saved for analysis to determine protein enrichment and capture by Western blot. Eight M urea (32 µL) was added and chymotrypsin (2 µg) was added to the beads and digestion was allowed to occur for 16 h at 24 °C. Beads were pelleted, and supernatant was transferred to a new tube. Beads were washed 3× with 1× PBS (200 µL) and the washes were transferred to the supernatant tube. Sample was concentrated to dryness using a vacuum centrifuge (i.e., a speedvac). Samples were desalted with a ZipTip P10, concentrated to dryness, and stored at –20 °C until analysis.

### PEG-5kDa glycoprotein labeling

Mass shift assays were performed according to the procedure of Pratt and co-workers (42). Briefly, samples (200 µg) were reduced with 25 mM DTT and heated for 5 min at 95 °C. Samples were then alkylated with 50 mM iodoacetamide for 1 h in the dark at 24 °C. Samples were precipitated by the addition of methanol (600 µL), chloroform (200 µL), and water (450 µL), vortexing, and centrifugation (10 min, 10,000 × g). The aqueous upper layer was discarded and methanol (1 mL) was added, sample was vortexed, and centrifuged (10 min, 10,000 × g). Sample was allowed to air dry before resuspension in 2% SDS + 1× PBS (45 µL) by probe tip sonication. Ten mM DBCO-PEG5K (5 µL, Click Chemistry Tools) was added and the solution warmed in a heat block for 5 min at 95 °C. Samples were precipitated by the addition of methanol (600 µL), chloroform (200 µL), and water (450 µL), vortexing and centrifugation (10 min, 10,000 × g). Aqueous upper layer was discarded and methanol (1 mL) was added, sample was vortexed, and centrifuged (10 min, 10,000 × g). Sample was allowed to air dry before resuspension by probe tip sonication in 2% SDS + 1× PBS (40 µL). 5× Laemmli sample buffer (10 µL) was added and the samples were heated for 5 min at 95 °C for Western blot analysis.

### Chemoenzymatic labeling of O-GlcNAc by Y289L GalT1 enzyme and chemical enrichment

Y289L GalT1 enzyme was expressed and purified following the procedure of Hsieh-Wilson and co-workers (43). Briefly, 2 mg of cell lysates (400 µL), which had been previously reduced and alkylated, were mixed with water (490 µL), GalT1 labeling buffer (800 μL, final concentrations: 50 mM NaCl, 20 mM HEPES, 2% NP-40, pH 7.9), and 100 mM MnCl_2_ (110 μL) were added in order. The sample was vortexed and transferred to ice. Then, 500 μM UDP-GalNAz (100 μL) and 2 mg/mL GalT1 enzyme (100 μL) were added to the sample. Subsequently, the sample reaction was rotated for 16 h at 4 °C. Samples were precipitated by the addition of methanol (1.2 mL), chloroform (400 µL), and water (900 µL), vortexing and centrifugation (10 min, 10,000 × g). Aqueous upper layer was discarded and methanol (1 mL) was added, sample was vortexed, and centrifuged (10 min, 10,000 × g). Sample was allowed to air dry before resuspension in 2% SDS + 1× PBS (400 µL). A pre-mixed solution of the click chemistry reagents (100 µL; final concentration of 200 µM IsoTaG silane probe, 500 µM CuSO_4_, 100 µM THPTA, 2.5 mM sodium ascorbate) was added and the reaction was incubated for 3.5 h at 24 °C. Samples were precipitated by the addition of methanol (600 µL), chloroform (200 µL), and water (450 µL), vortexing and centrifugation (10 min, 10,000 × g). Aqueous upper layer was discarded and methanol (1 mL) was added, sample was vortexed, and centrifuged (10 min, 10,000 × g). Sample was allowed to air dry before resuspension in 2% SDS + 1× PBS (400 µL) by probe tip sonication. Streptavidin–agarose resin [400 µL of the resin slurry, washed with PBS (3 × 1 mL)] was added, and the resulting mixture was incubated for 12 h at 24 °C with rotation. The beads were washed using spin columns with 8 M urea (5 × 1 mL), and PBS (5 × 1 mL). Washed beads were resuspended in 1× PBS + 10 mM CaCl_2_ (520 µL). Fifty µL of this mixture was saved for analysis to determine protein enrichment and capture by Western blot. Eight M urea (32 µL) and trypsin (1.5 µg) was added to the beads and digestion was allowed to occur for 16 h at 37 °C with rotation. Supernatant was collected from spin columns and the beads were washed with PBS (1 × 200 µL) and H_2_O (2 × 200 µL). Washes were combined with the supernatant digest to form the trypsin digest. The IsoTaG silane probe was cleaved with 2% formic acid/water (2 × 200 µL) for 30 min at 24 °C with rotation and the eluent was collected. The beads were washed with 50% acetonitrile– water + 1% formic acid (2 × 500 µL), and the washes were combined with the eluent to form the cleavage fraction. The trypsin digest and cleavage fraction were concentrated using a vacuum centrifuge (i.e., a speedvac) to dryness and then resuspended with 2% formic acid/water (50 µL). Samples were desalted with a ZipTip P10. Trypsin fractions were resuspended in 50 mM TEAB (20 µL) and TMT reagent (2 µL) was added to the samples and incubated for 1 h at 24°C. Hydroxyammonia (50%, 1 µL) was added to the samples to quench the reaction for 15 min at 24°C. Samples were combined and concentrated using a vacuum centrifuge (i.e., a speedvac) to dryness and stored at –20 °C until analysis.

### Western blot procedures

The protein sample (15 µL) was loaded on 6–12% or 6–10% Tris-Glycine SDS-PAGE gels and ran on a Mini-PROTEAN® BioRad gel system. Gels were transferred with the Invitrogen iBlot. For α-synuclein blots, membranes were incubated in 1% paraformaldehyde for 1 h to prevent α-synuclein dissociation from the membrane as previously described prior to blocking (44). Membranes were stained with LI-COR Revert total protein stain to verify transfer and equal protein loading and blocked with 3% BSA + 1× TBST for 1 h at 24 °C. Primary antibodies and the following dilutions were incubated with the membranes for 1 h to 12 h: anti-Flag (1:5,000; Sigma Aldrich; Cat # F3165), anti-Flag (1:1,000; Cell Signaling; Cat # 14793S), anti-HA (1:1,000; Cell Signaling; Cat # 3724S), anti-O-GlcNAc RL2 (1:1,000; Abcam; Cat # ab2739), anti-synuclein (1:1,000; Abcam; Cat # ab138501). Membranes were washed 3 × 5 min each wash with 1× TBST and incubated with the following secondary antibodies and dilutions: anti-Mouse HRP (1:10,000; Rockland Immunochemicals: Cat # 610-1302), anti-Rabbit HRP (1:10,000; Rockland Immunochemicals: Cat # 611-1302), anti-Mouse IR 800 (1:10,000; LI-COR; Cat # 925-32210), anti-Rabbit IR 680 (1:10,000; LI-COR; Cat # 925-68071), anti-Rabbit IR 800 (1:10,000; LI-COR; Cat # 925-32211). Membranes were washed 3 × 5 min each wash with 1× TBST and results obtained by chemiluminescence or IR imaging using the Azure c600. Membranes were quantified using LI-COR image studio lite.

### Mass spectrometry acquisition procedures(15)

Desalted samples were reconstituted in 0.1% formic acid in water (20 µL), and half of the sample (10 µL) was injected onto a C18 trap column (WATERS Cat # 186008821 nanoEase MZ Symmetry C18 Trap Column, 100 Å, 5 µm × 180 µm × 20 mm) and separated on an analytical column (WATERS Cat # 186008795 nanoEase MZ Peptide BEH C18 Column, 130 Å, 1.7 µm × 75 µm × 250 mm) with a Waters nanoAcquity system connected in line to a ThermoScientific Orbitrap Fusion Tribrid. The column temperature was maintained at 50 °C. Peptides were eluted using a multi-step gradient at a flow rate of 0.15 µL/min over 120 min (0−5 min, 2−5% acetonitrile in 0.1% formic acid in water; 5−95 min, 5−50%; 95−105 min, 50−98%; 105−115 min, 98%; 115– 116 min, 98–2%; 116–120 min, 2%). The electrospray ionization voltage was set to 2 kV and the capillary temperature was set to 275 °C. Dynamic exclusion was enabled with a repeat count of 2, repeat duration of 30 s, exclusion list size of 400, and exclusion duration of 30 s. MS1 scans were performed over 400–2000 *m/z* at resolution 120,000 and the top twenty most intense ions (+2 to +6 charge states) were subjected to MS2 HCD fragmentation at 27%, for 75 ms, at resolution 50,000. Other relevant parameters of HCD include: isolation window (3 *m/z*), first mass (100 *m/z*), and inject ions for all available parallelizable time (True). If oxonium product ions (138.0545, 204.0867, 345.1400, 347.1530 *m/z*) were observed in the HCD spectra, ETD (250 ms) with supplemental activation (35%) was performed in a subsequent scan on the same precursor ion selected for HCD. Other relevant parameters of ETHCD include: isolation window (3 *m/z*), use calibrated charge-dependent ETD parameters (True), Orbitrap resolution (50k), first mass (100 *m/z*), and inject ions for all available parallelizable time (True).

### Mass spectrometry data analysis

The raw data was processed using Proteome Discoverer 2.3 (Thermo Fisher Scientific). For quantitative proteomics, the data was searched against the human-specific SwissProt-reviewed database 2016 (20,152 proteins, downloaded on Aug. 19, 2016). HCD spectra with a signal-to-noise ratio greater than 1.5 were searched against a database containing the Swissprot 2016 annotated human proteome and contaminant proteins using Sequest HT with a mass tolerance of 10 ppm for the precursor and 0.02 Da for fragment ions with specific trypsin digestion, 2 missed cleavages, variable oxidation on methionine residues (+15.995 Da), static carboxyamidomethylation of cysteine residues (+57.021 Da), and static TMT labeling (229.163 Da) at lysine residues and peptide N-termini. High confidence assignments were filtered using Percolator (FDR = 1%). The TMT reporter ions were quantified using the Reporter Ions Quantifier and normalized such that the summed peptide intensity per channel was equal.

For all glycoproteomics data, the data was searched using Byonic v3.0.0 as a node in Proteome Discoverer 2.3 against the target protein sequence (Nup62, P37198; JunB, P17275), chymotrypsin, and the nEPEA(4) construct. Indexed databases for chymotryptic digests were created with full cleavage specificity. The database allowed for up to three missed cleavages with variable modifications (methionine oxidation, +15.9949 Da; carbamidomethylcysteine, +57.0215 Da; deamidation of asparagine and glutamine, +0.984016 Da; and others as described below). Precursor ion mass tolerances for spectra acquired using the Orbitrap were set to 10 ppm. Glycopeptide searches allowed for tagged HexNAc modifications (HexNAc +203.0794 on serine, threonine). Glycopeptide spectral assignments passing a FDR of 1% at the peptide spectrum match level based on a target decoy database were kept. Singly modified glycopeptides assigned from EThCD spectra passing a 1% FDR and possessing a delta modification score of greater than or equal to ten from >2 PSMs were considered unambiguous glycosites. All other glycopeptides passing a 1% FDR were considered ambiguous glycosites.

### Immunofluorescence and fixed-cell sample preparation

Cells were seeded on 22×22 mm glass coverslips no. 1.5 coated with poly-L-lysine (Neuvitro Corporation German Glass Coverslips Cat # H-22-1.5-pll) that had been placed in single wells of a 6-well plate for 24 h prior to transfection. For experiments in Figure 5A, cells were plated in normal-glucose (1 g/L) DMEM (Corning, Cat # 10014CV). Cells were transfected for 48 h and the media were exchanged with fresh media after 24 h. For experiments in Figure 5A, cells were transfected in a 6-well plate without coverslips for 24 h, trypsinized, and replated on a glass coverslip mentioned above. The replated cells were kept for an additional 24 h. Transfected cells were fixed in freshly prepared 4% paraformaldehyde in PBS (pH 7.4) for 15 min at 24 °C (1 mL per well), washed with PBS (2 mL per well) twice (10 min), permeabilized in PBS with 0.1% Triton X-100 (1 mL per well) for 20 min at 24 °C. Cells were washed with PBS for 15 min (3 × 2 mL), and then incubated with blocking solution (3% BSA/TBST) for at least 1 h at 4 °C, followed by overnight incubation with the primary antibody. Cells were washed with PBS for 15 min (3 × 2 mL), and subsequently incubated with the secondary antibody for 1 h at 4 °C, washed with PBS for 15 min (3 × 2 mL). The nuclei were stained in DAPI solution (4′,6-diamidino-2-phenylindole, Invitrogen Molecular Probes NucBlue, Cat # R37606) for 10 min at 24 °C. Coverslips were mounted in anti-fade Diamond (Life Technologies Cat # P36961). Primary antibodies was rabbit anti-HA mAb (1:1000, HA-Tag-C29F4 Cell Signaling, Cat # 3724S). Secondary antibodies was goat anti-Rabbit IgG (H+L) cross-adsorbed secondary antibody conjugated to Alexa Fluor 594 (1:5000, ThermoFisher/Invitrogen, Cat # A11012).

### Confocal fluorescence microscopy, image acquisition and processing

Fixed-cell samples were imaged using a Zeiss laser scanning confocal microscope (LSM) 880 confocal microscope. Images were acquired with a Plan-Apochromat 40X or 63X/1.4NA oil immersion objective DIC M27 (the magnification was adjusted by zooming in or out as needed). Excitation wavelengths for DAPI, Alexa Fluor488, red fluorescent protein (RFP), and Alexa Fluor594 were at 405 nm, 488 nm, 561 nm, and 594 nm, respectively. The laser power and detector gain were adjusted to obtain the best signal-to-noise ratio and have no over-saturated signal. Fluorescence was detected using the Zeiss QUASAR detection unit. Sequential Z stacks were acquired consisting of 11 planes separated by 0.5 µm, pixel size 0.19 µm, with a 0.52 µs pixel dwell time (2×2 averaging per frame was used). A pinhole size of 1 Airy Unit (AU) at all wavelengths was used. Images were processed with ImageJ2 (Fiji). All images shown are average-intensity projections from all slices in z-stacks.

### Statistical analyses

Statistical analyses methods are described in figure legends. Two tailed t-tests and one-way ANOVA tests were performed.

