## supplementary information for "Selective protein O-GlcNAcylation in cells by a proximity-directed O-GlcNAc transferase"


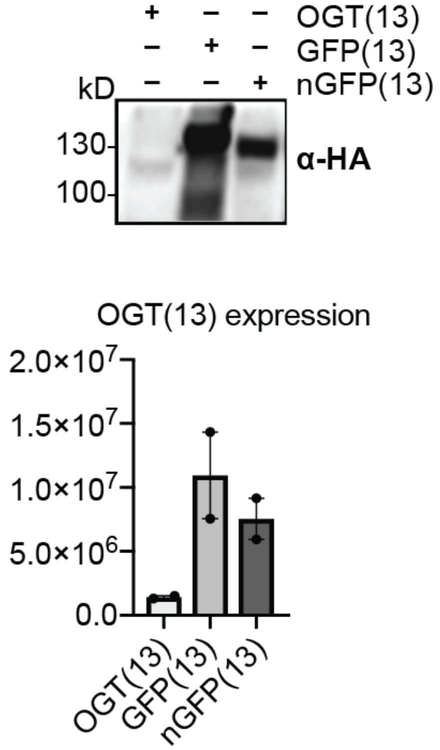


**Supplementary Figure S1 |** Western blot for expression of the indicated OGT construct. HEK293T cells were transfected with the indicated construct, cell lysates were collected, and a Western blot was performed for the HA-tag.

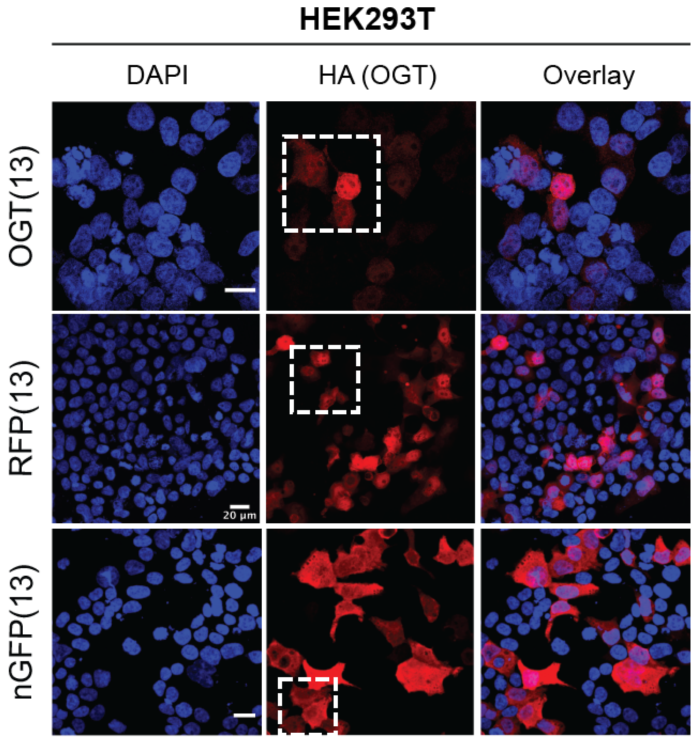


**Supplementary Figure S2 |** Wide field confocal imaging of Figure 2. Scale bars represent 20 μm.

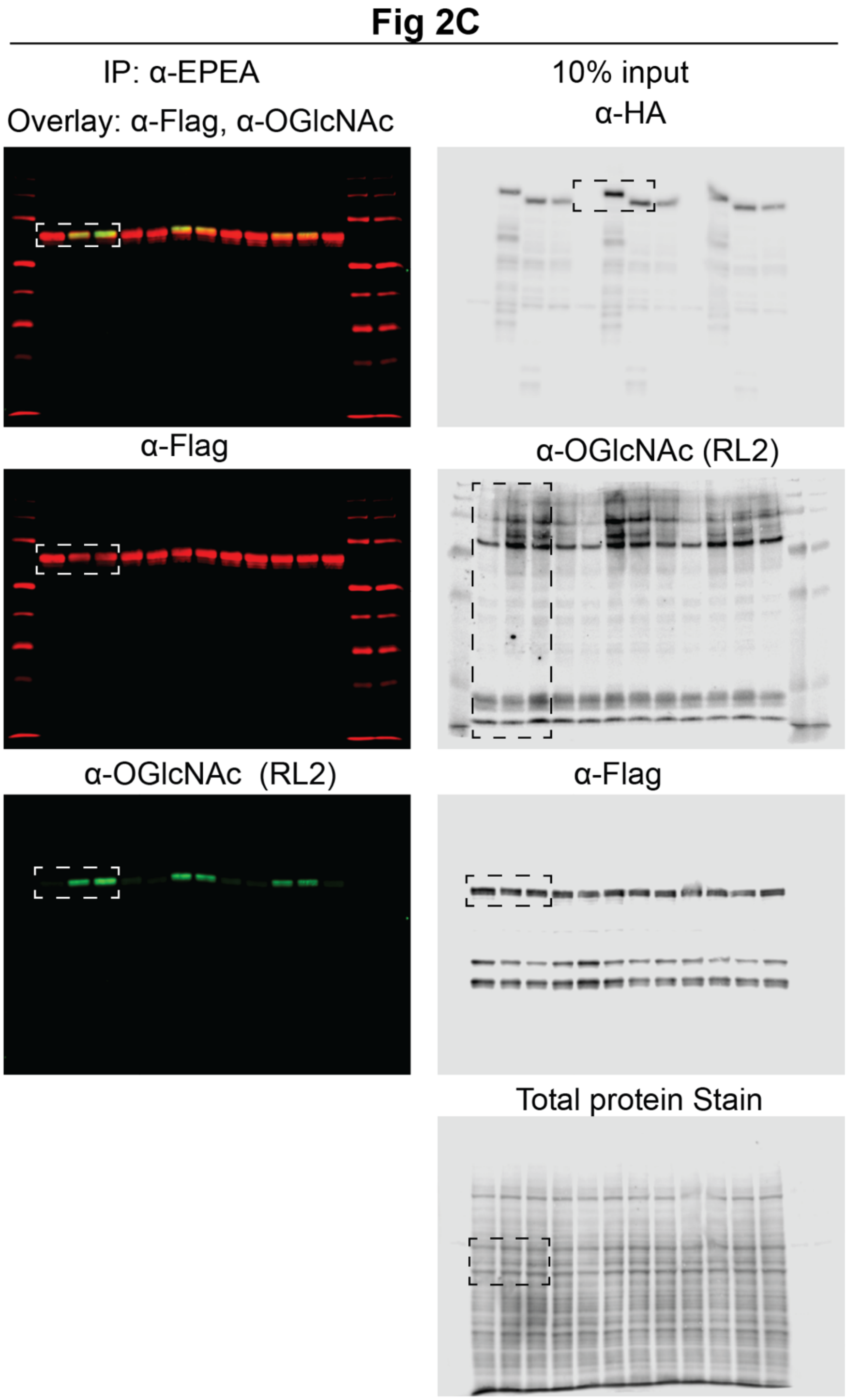


**Supplementary Figure S3 |** Uncropped Western blot and total protein stain images in Figure 2.

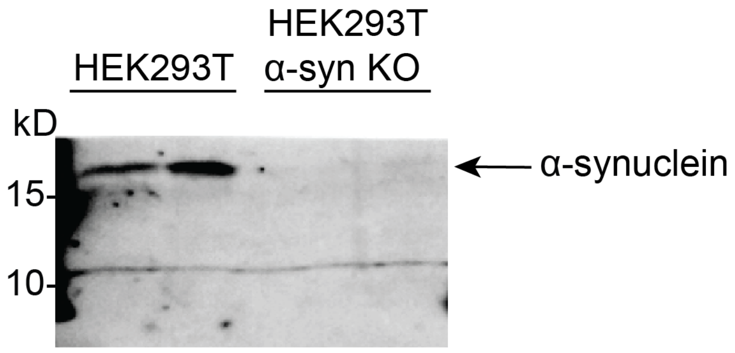


**Supplementary Figure S4 |** Western blot for α-synuclein in HEK293T and α-syn KO HEK293T cells.

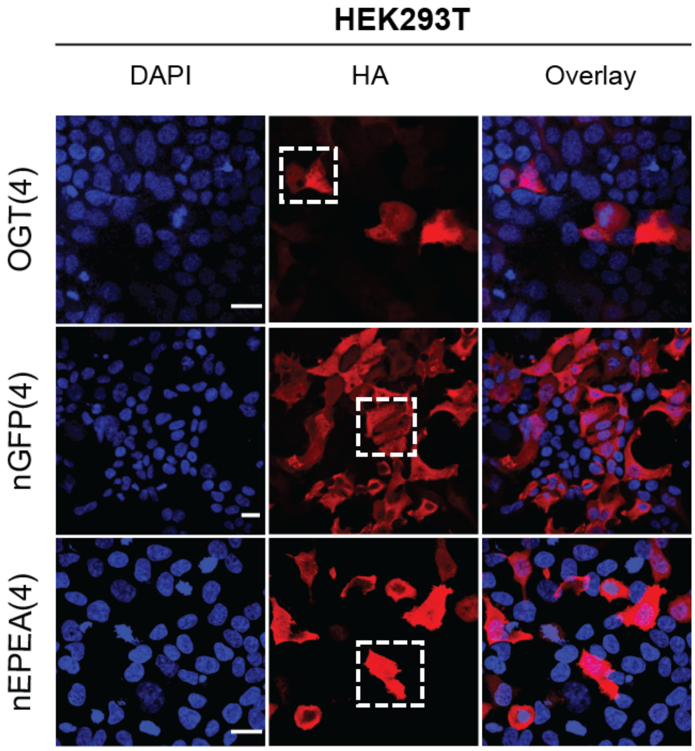


**Supplementary Figure S5 |** Wide field confocal imaging of Figure 3. Scale bars represent 20 μm.

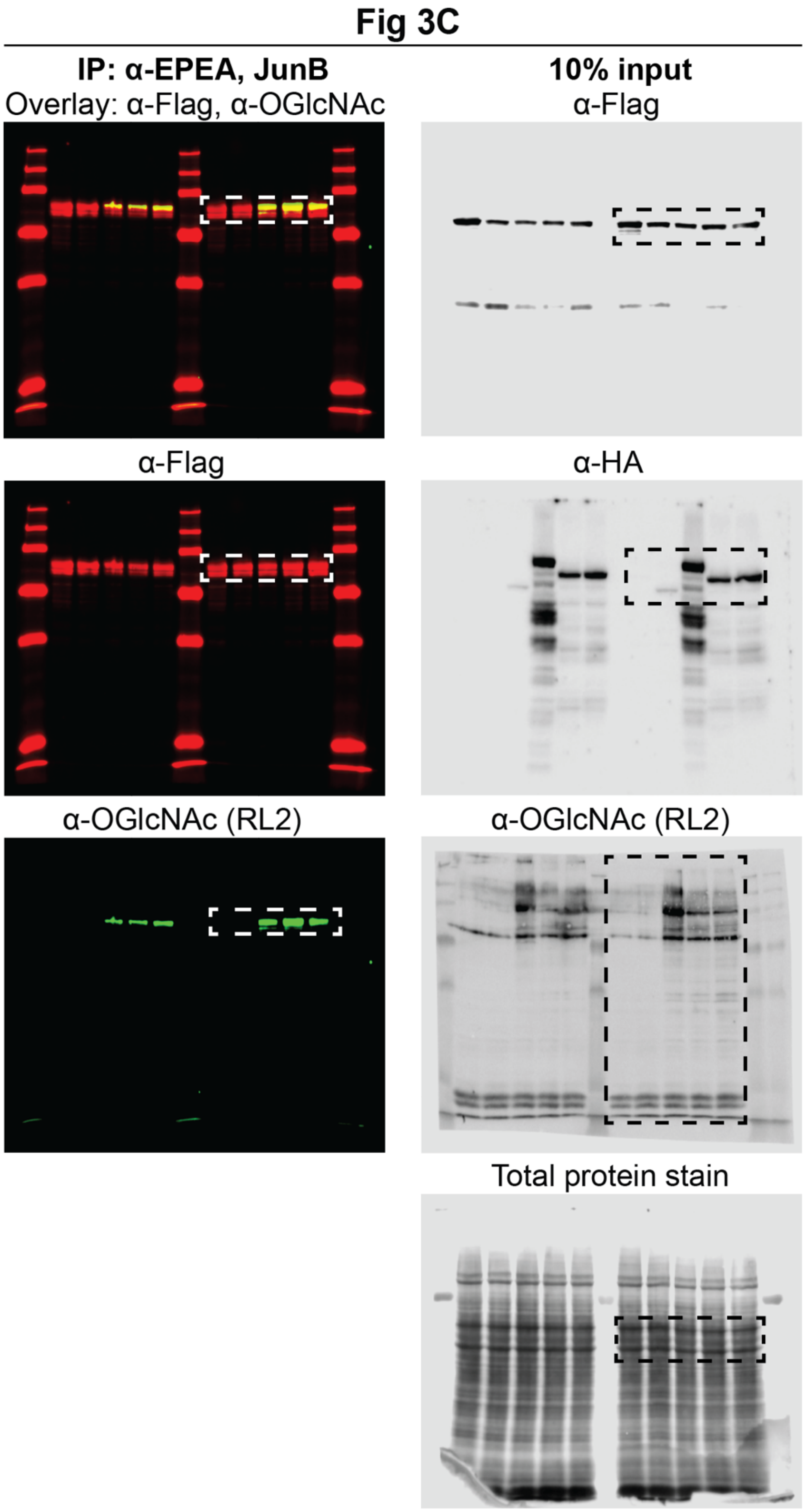


**Supplementary Figure S6 |** Uncropped Western blot and total protein stain images in Figures 3C

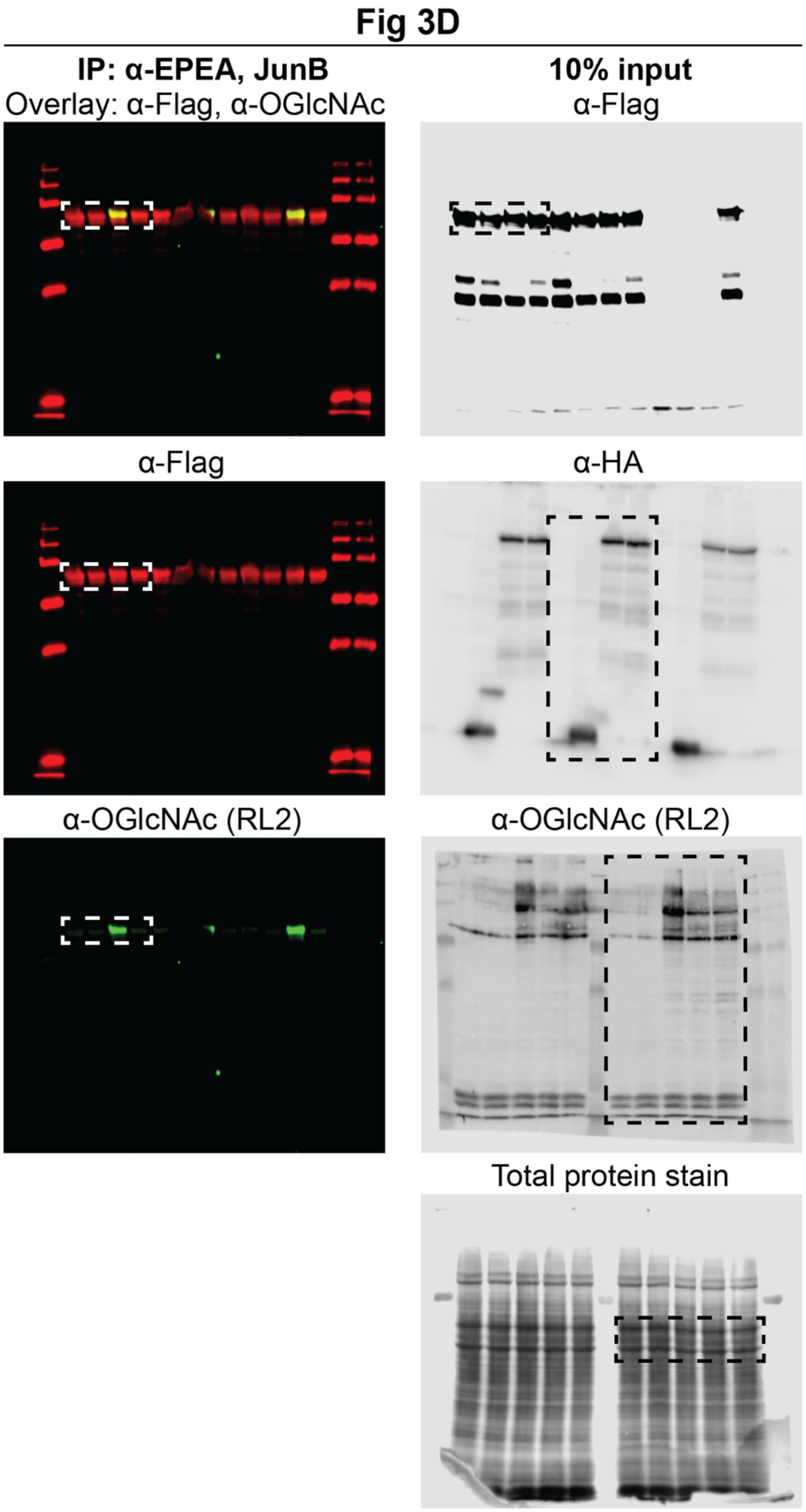


**Supplementary Figure S7 |** Uncropped Western blot and total protein stain images in Figures 3D

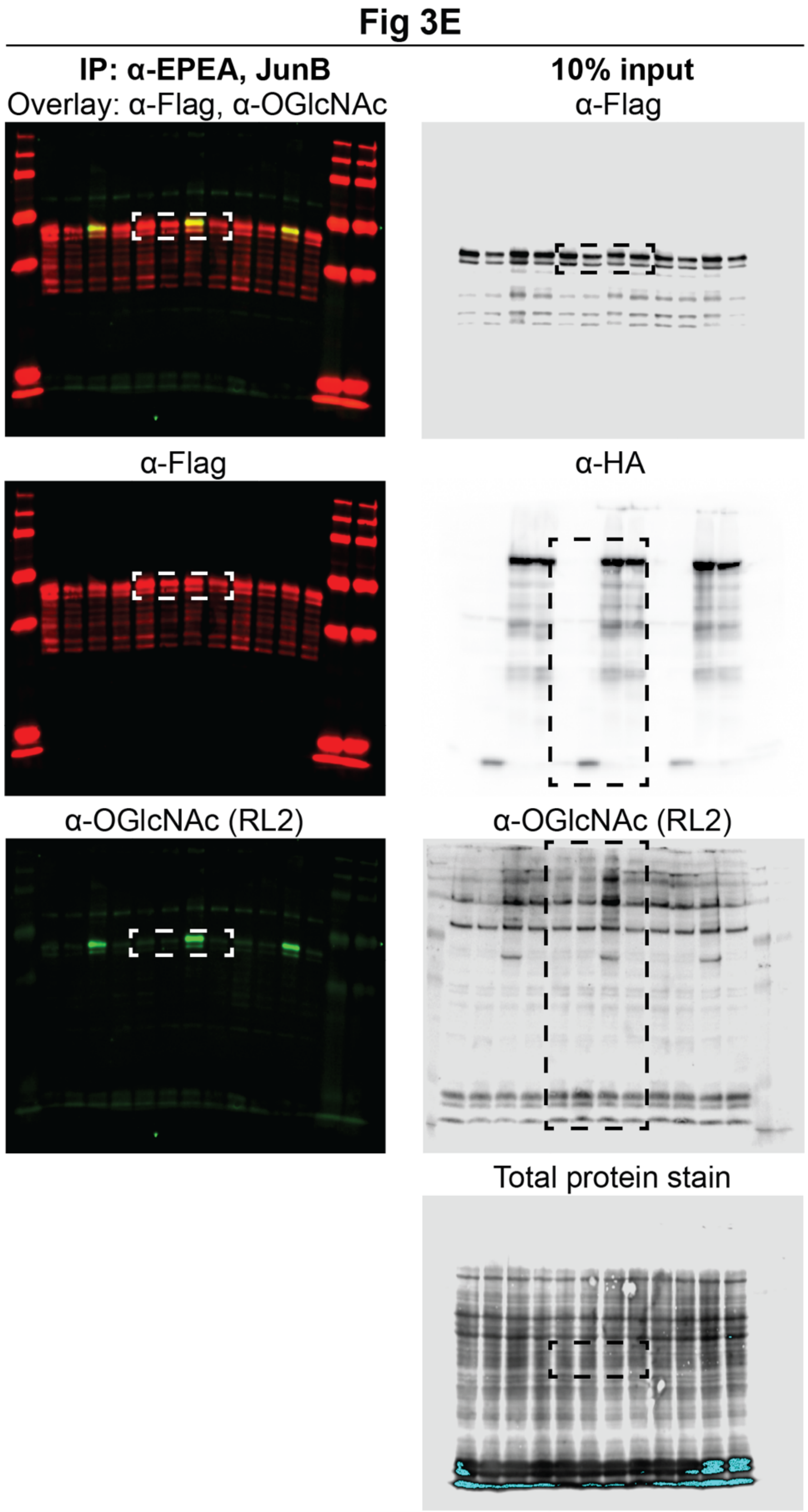


**Supplementary Figure S8 |** Uncropped Western blot and total protein stain images in Figures 3E

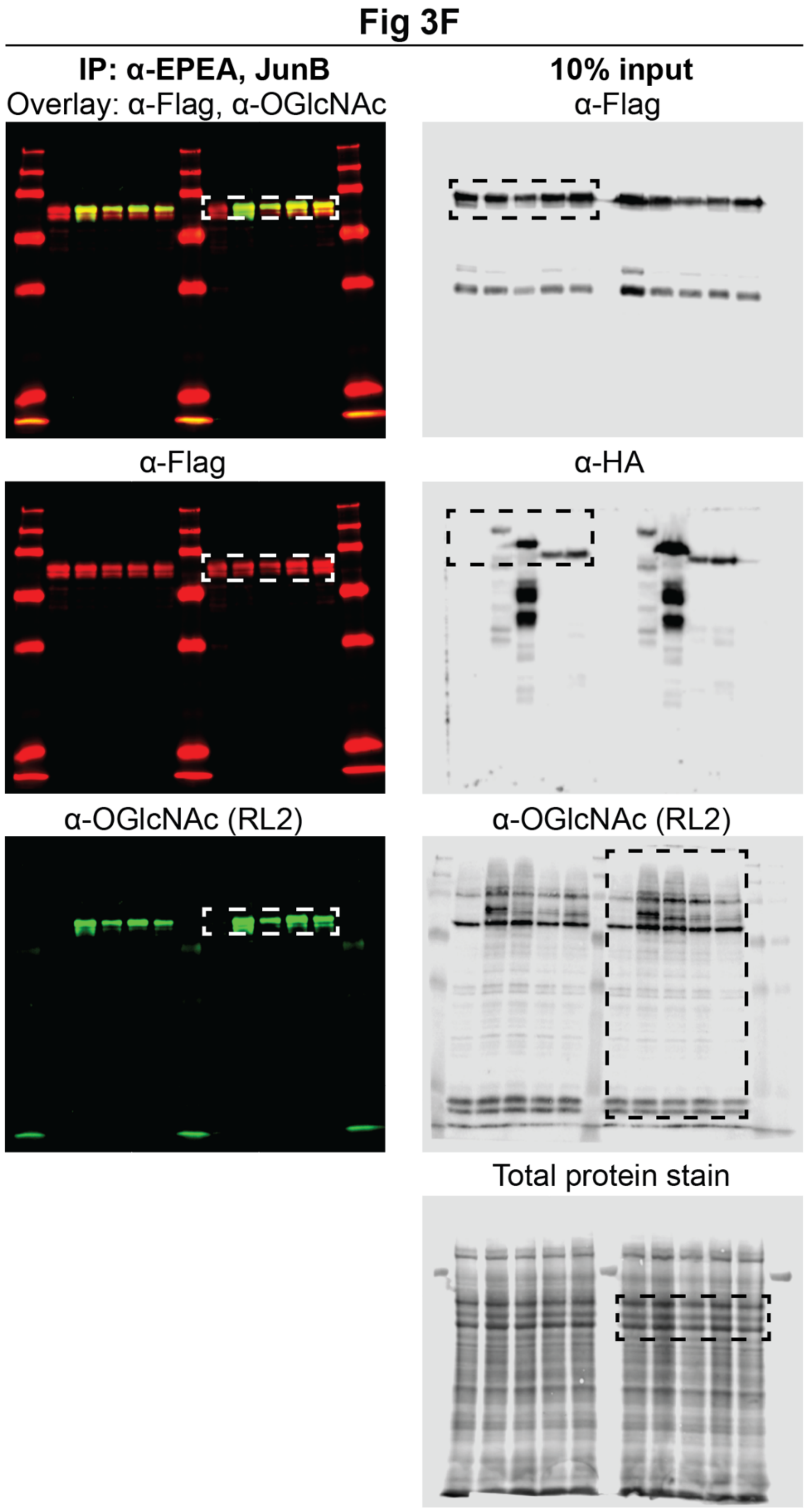


**Supplementary Figure S9 |** Uncropped Western blot and total protein stain images in Figures 3F

### Fig 3G

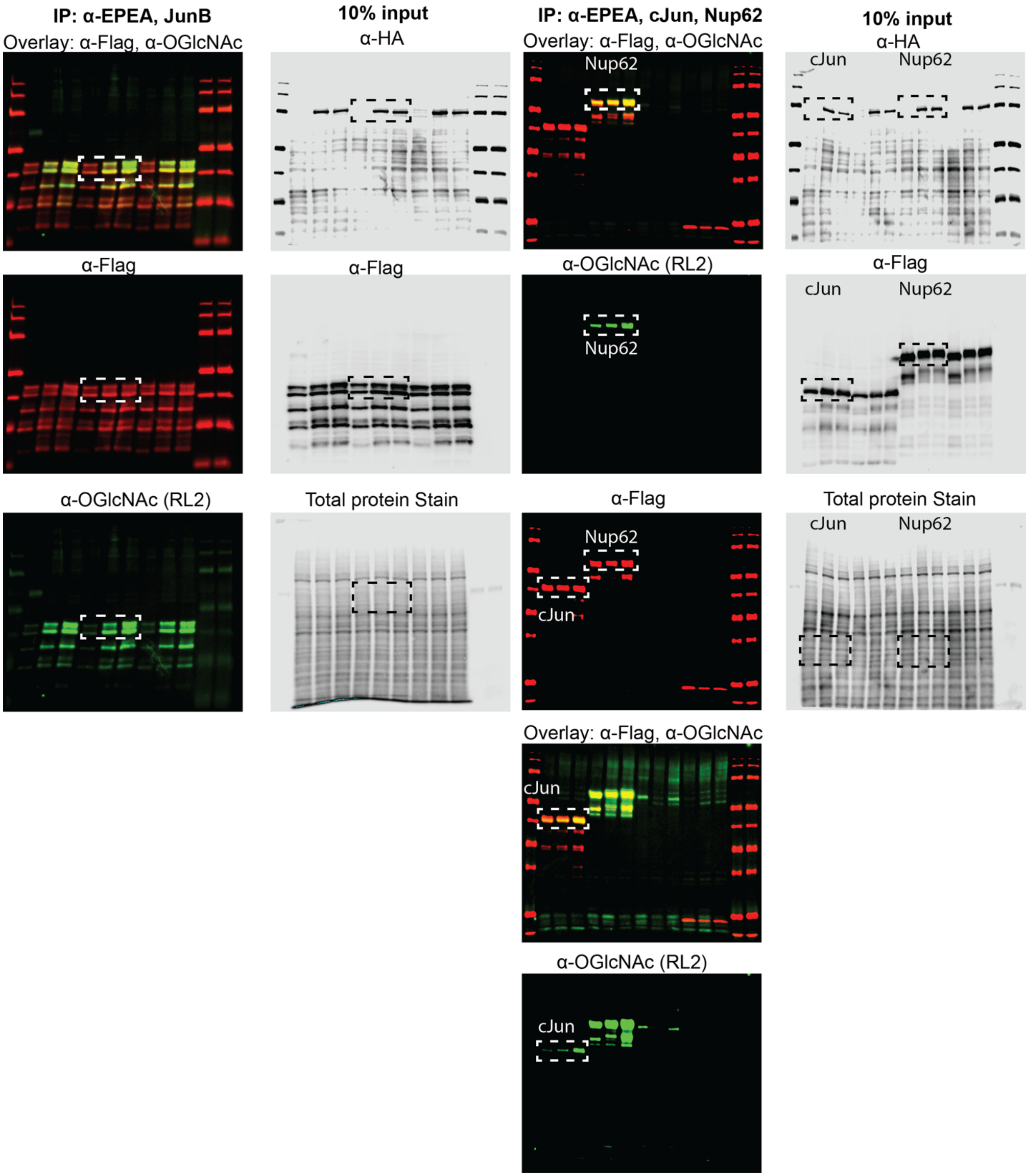


**Supplementary Figure S10 |** Uncropped Western blot and total protein stain images in figures 3G

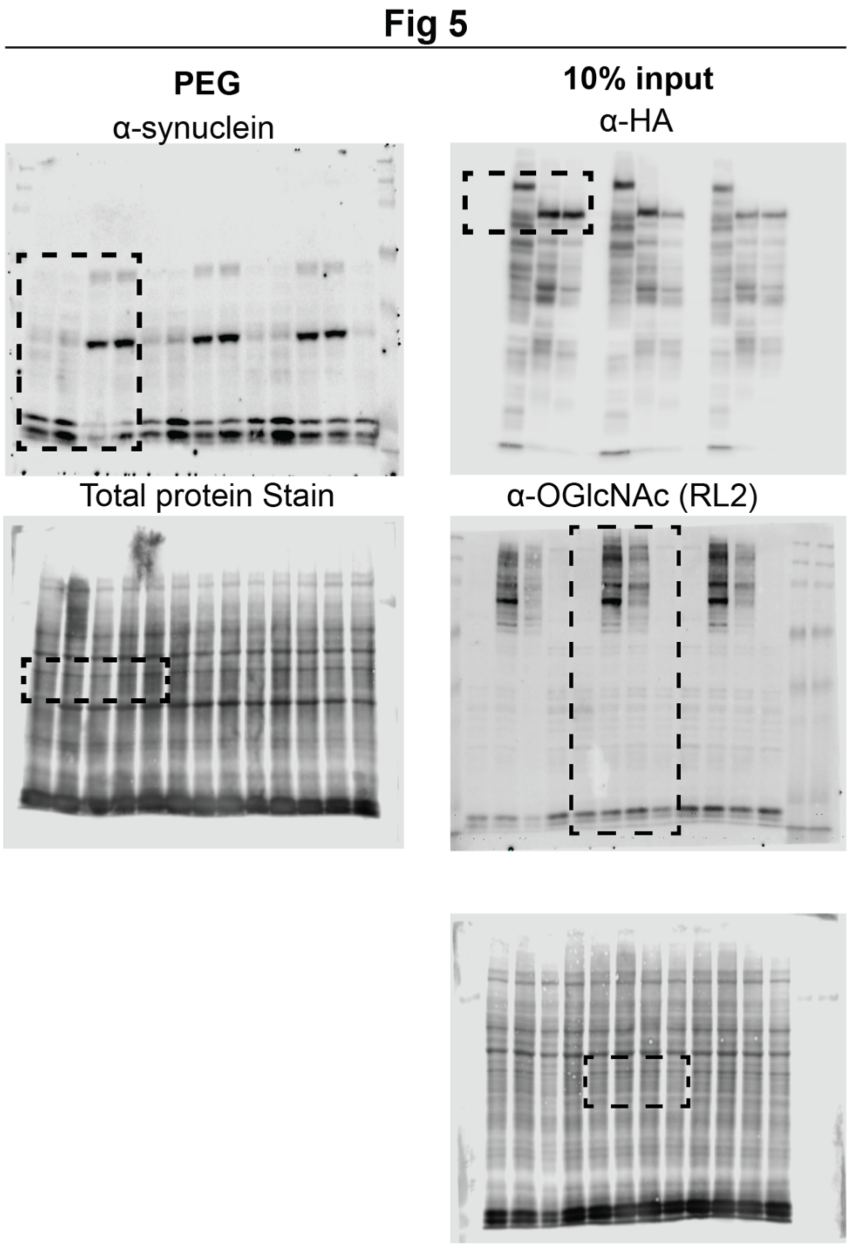


**Supplementary Figure S11 |** Uncropped Western blot and total protein stain images in Figure 5.

**Supplementary Information.**

**Abbreviations**

| | |
|---|---|
| α-syn KO | alpha-synuclein knockout |
| BSA | Bovine Serum Albumin |
| CID | Collision Induced Dissociation |
| CuAAC | Copper-catalyzed azide-alkyne cyclooaddition |
| DAPI | (4′,6-diamidino-2-phenylindole) |
| DBCO-PEG5K | Dibenzocyclooctyne-Polyethylene glycol 5kDa |
| DMEM | Dulbecco's Modified Eagle's medium |
| DMSO | Dimethyl sulfoxide |
| EDTA | Ethylenediaminetetraacetic acid |
| EThCD | Electron transfer higher-energy collision dissociation |
| FBS | Fetal Bovine Serum |
| GALE | UDP-Galactose-4-epimerase |
| GFP | Green Fluorescent Protein |
| HCD | High collision dissociation |
| HDR | Homology directed repair |
| HexNAc | N-Acetyl hexosamine |
| HexNAc0Si | IsoTaG-clicked N-azidoacetyl hexosamine |
| HexNAc2Si | IsoTaG-clicked N-azidoacetyl hexosamine |
| IDT | Integrated DNA technologies |
| IsoTaG | Isotope targeted glycoproteomics |
| KO | Knockout |
| MS | Mass Spectrometry |
| nEPEA | EPEA nanobody |
| nGFP | GFP nanobody |
| O-GlcNAc | O-linked-ß-N-Acetylglucosamine |
| OGA | O-GlcNAcase |
| OGT | O-GlcNAc transferase |
| OGT(13) | OGT with 13.5 TPRs |
| OGT(4) | OGT with 4.5 TPRs |
| PEG | Polyethylene glycol |
| PSM | Peptide spectral match |
| PTM | Post-translational modification |
| RE | Restriction Enzyme |
| TBST | Tris-buffered saline + 0.1% Tween |
| THPTA | Tris(3-hydroxypropyltriazolylmethyl)amine |
| TPR | Tetratricopeptide repeat |
| UDP-GalNAz | Uridine diphosphate N-azidoacetyl galactosamine |

**Synthesized materials and storage**

1. The cleavable biotin silane probe as a 1:3 ratio mixture of the light and heavy (+2 deuteriums) stable isotopes was prepared according to the procedure of Bertozzi and co-workers.[1] The cleavable biotin silane probe was dissolved in DMSO to obtain a 10 mM stock solution and kept in amber microcentrifuge tubes at −20 °C for short-term storage and kept as in solid form at −80 °C for long-term storage.

**Purchased reagents**

| No. | Reagent name | Source | Catalog # |
|---|---|---|---|
| 1 | 150-mm cell culture dishes | Corning | 25383-103 |
| 2 | 6-well plate | VWR | 10062-892 |
| 3 | Anti-Fade Diamond | Life Technologies | P36961 |
| 4 | BSA | Sigma-Aldrich | A7906-1KG |
| 5 | C-tag resin | Thermo Fisher | 191307005 |
| 6 | C-tag resin XL | Thermo Fisher | 2943072005 |
| 7 | Chymotrypsin, Sequencing grade | Promega | V1062 |
| 8 | cOmplete™, EDTA-free Protease Inhibitor Cocktail | Sigma Aldrich | 11873580001 |
| 9 | DBCO–PEG-5kDa | Click Chemistry Tools | A118-100 |
| 10 | Dulbecco's Modified Eagle Medium | Lonza | 12-604F |
| 11 | FBS | VWR Life Science | 97068-085 |
| 12 | Fetal Bovine Serum (FBS) | VWR | 97068-085 |
| 13 | German Glass Coverslips | NeuVitro | H-22-1.5-pll |
| 14 | iBlot Transfer Stack, nitrocellulose | Thermo Fisher | IB301001 |
| 15 | LI-COR REVERT Total Protein Stain | LI-COR Biosciences | 926-11016 |
| 16 | Lipofectamine 2000 | Thermo Fisher | 11668019 |
| 17 | Mini Bio-Spin Chromatography columns | Bio-Rad | 7326207 |
| 18 | NucBlue | Invitrogen | R37606 |
| 19 | Opti-MEM | Thermo Fisher | 31985070 |
| 20 | Penicillin-Streptomycin | Life Technologies | 15140122 |
| 21 | Pierce™ TMT10plex™ Isobaric Mass Tag Labelling Kit and | Thermo Fisher | 90406 |
| 22 | Streptavidin-Agarose | Thermo Fisher | 20361 |
| 23 | Thiamet-G | Sigma-Aldrich | SML0244-25MG |
| 24 | THPTA | Sigma-Aldrich | 762342 |
| 25 | Transit-Pro | Mirus Bio | MIR 5740 |
| 26 | Trypsin-EDTA (0.25%), phenol red | Thermo Fisher | 25200114 |
| 27 | Trypsin, Sequencing grade | Promega | V5111 |
| 28 | ZipTip P10 | Millipore Sigma | ZTC18S096 |

**Antibodies**

All antibodies were diluted in 3% BSA/TBST, unless otherwise noted.

| No. | Antibody name | Host species | Dilution | Commercial source | Catalog # |
|---|---|---|---|---|---|
| 1 | Flag (M2) | Mouse mAb | 1:5,000 | Sigma-Aldrich | F3165 |
| 2 | HA-Tag (C29F4) | Rabbit mAb | 1:1,000 | Cell Signaling | 3724S |
| 3 | O-GlcNAc (RL2) | Mouse mAb | 1:1,000 | Abcam | Ab2739 |
| 4 | Alpha-synuclein | Rabbit mAb | 1:1,000 | Abcam | Ab138501 |

| | | | | | |
|---|---|---|---|---|---|
| **5** | Flag (D6W5B) | Rabbit | 1:1,000 | Cell signaling | 14793S |
| **6** | Alexa Fluor 594 | Goat | 1:5,000 | Thermo Fisher/Invitrogen | A11012 |
| **7** | Anti-mouse-HRP | Goat | 1:10,000 | Rockland Immunochemicals | 610-1302 |
| **8** | Anti-rabbit-HRP | Goat | 1:10,000 | Rockland Immunochemicals | 611-1302 |
| **9** | Anti-mouse-IR 800 | Goat | 1:10,000 | LI-COR Biosciences | 925-32210 |
| **10** | Anti-rabbit-IR 680 | Goat | 1:10,000 | LI-COR Biosciences | 925-68071 |
| **11** | Anti-rabbit-IR 800 | Goat | 1:10,000 | LI-COR Biosciences | 925-32211 |

**Molecular Cloning Reagents**

| No. | Antibody name | Commercial source | Catalog # |
|---|---|---|---|
| **1** | Gibson Assembly Master mix | New England Biolabs | E2611L |
| **2** | Q5 High-Fidelity 2X Master Mix | New England Biolabs | M0492S |
| **3** | T4 Polynucleotide Kinase | New England Biolabs | M0201L |
| **4** | HindIII-HF | New England Biolabs | R3104M |
| **5** | NotI-HF | New England Biolabs | M0492S |
| **6** | SgsI | New England Biolabs | R0558L |
| **7** | SgfI | New England Biolabs | R0630L |
| **8** | BamHI-HF | New England Biolabs | R3136L |
| **9** | XhoI | New England Biolabs | R0146L |

**Cloning Plasmids**

| No. | Plasmid name | Source |
|---|---|---|
| **1** | pCSDEST2-APEX2-GBP | A gift from Rob Parton (Addgene Plasmid # 67651) |
| **2** | pEXP pLHCX ncOGT | A gift from the Pratt lab |
| **3** | pEGFP-(C3)-Nup153 | A gift from Birthe Fahrenkrog (Addgene plasmid # 64268) |
| **4** | pCS-H2B-mRFP | A gift from Sean Megason (Addgene plasmid # 53745) |
| **5** | JunB | A gift from the Davis lab |
| **6** | pDONR223-NUP62 | A gift from William Hahn & David Root (Addgene Plasmid # 23559) |
| **7** | cJun | A gift from the Davis lab |

**Plasmids**

All plasmids in this study are derived from the Invitrogen pcDNA3.1 vector, which contains a CMV promoter for constitutive expression.

| No. | Plasmid No. | Plasmid name |
|---|---|---|

| | | |
|---|---|---|
| **1** | pWLH085 | pcDNA3.1-HA-nGFP-(EAAAK)4-OGT (13) |
| **2** | pWLH216 | pcDNA3.1-HA-GFP-(EAAAK)4-OGT(13) |
| **3** | pWLH300 | pcDNA3.1-HA-RFP-(EAAAK)4-OGT(13) |
| **4** | pWLH118 | pcDNA3.1-HA-OGT (13) |
| **5** | pWLH117 | pcDNA3.1-HA-OGT(4) |
| **6** | pWLH259 | pcDNA3.1-HA-GFP-(EAAAK)4-OGT(4) |
| **7** | pWLH189 | pcDNA3.1-HA-nGFP-(EAAAK)4-OGT(4) |
| **8** | pWLH137 | pcDNA3.1-HA-nEPEA-(EAAAK)4-OGT(4) |
| **9** | pWLH282 | pcDNA3.1-HA-nGFP-(EAAAK)4-OGT(4,K852A) |
| **10** | pWLH194 | pcDNA3.1-HA-nEPEA-(EAAAK)4-OGT(4,K852A) |
| **11** | pWLH114 | pcDNA3.1-GFP-Flag-JunB-EPEA |
| **12** | pWLH147 | pcDNA3.1-Nup62-Flag-EPEA |
| **13** | pWLH142 | pcDNA3.1-cJun-Flag-EPEA |
| **14** | pWLH082 | pcDNA3.1-JunB-Flag-EPEA |

**Gene Blocks**

| | **Sequence** |
|---|---|
| **EPEA nanobody** | ATGGGCCAGCTGGTGGAGAGCGGCGGCGGCAGCGTGCAGGCCGGCGGCAGCCTGAGGCTGAGCTGCGCCGCCAGCGGCATCGACAGCAGCAGCTACTGCATGGGCTGGTTCAGGCAGAGGCCCGGCAAGGAGAGGGAGGGCGTGGCCAGGATCAACGGCCTGGGCGGCGTGAAGACCGCCTACGCCGACAGCGTGAAGGACAGGTTCACCATCAGCAGGGACAACGCCGAGAACACCGTGTACCTGCAGATGAACAGCCTGAAGCCCGAGGACACCGCCATCTACTACTGCGCCGCCAAGTTCAGCCCCGGCTACTGCGGCGGCAGCTGGAGCAACTTCGGCTACTGGGGCCAGGGCACCCAGGTTACTGTGAGCTCTCACCACCATCATCATCATCTGCCCGAGACCGGC |

**Primers**

| **No** | **Primer name** | **Sequence (5' to 3')** |
|---|---|---|
| **1** | pcDNA3.1-HindIII-HA-SgfI-nGFP fwd | CCCAAGCTGGCGAGCGTTTAAGCTTGAGCAATGGCATACCCATACGATGTTCCAGATTACGCTGCGATCGCACAGGTGCAGCTGGTGGAGTCTGGAGGA |
| **2** | (EAAAK)4-SgsI-nGFP rev | GGATCCCTTTGCAGCTGCCTCCTTTGCAGCTGCCTCCTTTGCAGCTGCCTCCTTTGCAGCTGCCTCTGGCGCGCCAGAGCTCACTGTCACCTGTGTT |
| **3** | (EAAAK)4-BamHI-OGT(1-1046) fwd | AAAGGAGGCAGCTGCAAAGGAGGCAGCTGCAAAGGGATCCATGGCGTCTTCCGTGGGCAA |
| **4** | pcDNA3.1-NotI-OGT(1-1046) rev | CGGGTTTAAACGGGCCCTCTAGACTCGAGCGGCCGCTTAGGCTGACTCGGTGACTTCAACAGGCTTAATCATGTGGTC |
| **5** | SgfI-GFP fwd | CGCTGCGATCGCAGTGAGCAAGGGCGAGGAGCTGTTCA |
| **6** | SgfI-RFP fwd | CGCTGCGATCGCAATGGCCTCCTCCGAGGACGTCATCAAGGAGTTC |
| **7** | SgsI-GFP rev | CTCTGGCGCGCCCTTGTACAGCTCGTCCATGCCGAGA |
| **8** | SgsI-RFP rev | TATTATTGGCGCGCCGGTGGAGTGGCGGCCCTCGGCGCGCTCGTACTGTT |

| | | |
|---|---|---|
| **9** | HindIII-HA-BamHI-OGT(1-1046) fwd | TTTAAGCTTGAGCAATGGCATACCCATACGATGTTCCAGATTACGCTGGATCCATGGCGTCTTCCGTGGGCAACGTGGCCGACAGTACA |
| **10** | NotI-OGT (1-1046) rev | TCGAGCGGCCGCTTAGGCTGACTCGGTGACTTCAACAGGCTTAATCATGTGGTCAGGTTT |
| **11** | BamHI-OGT(327–1046) fwd | AAAGGGATCCATGGCAGACTCTTTGAATAACCTTGCCAACATCAAACGGG |
| **12** | OGT(K852A) fwd | ATTGACCCATCTACCCTGCAGATGTGGGCAAATATTCTG |
| **13** | OGT(K852A) rev | TGCATATAACTGATTAAAGTTACAGTACACAATGGCATCTTCTGGTAGCCC |
| **14** | HindIII-GFP fwd | GCTAGCGTTTAAACTTAAGCTTGAGCAATGGTGAGCAAGGGCGAGG |
| **15** | (EAAAK)-Flag-SgsI-GFP rev | TTCCTTGGCGGCGGCCTCTCCCTTATCGTCGTCATCCTTGTAGTCTGGCGCGCCCTTGTACAGCTCGTCCATGCCGAGAGTGAT |
| **16** | (EAAAK)-Jun-B fwd | GGGGAGGCCGCCGCCAAGGAACGGATGTGCACTAAAATGGAACAGCCCTT |
| **17** | XhoI-EPEA-JunB rev | TTAAACGGGCCCTCTAGACTCGAGTTATGCTTCAGGTTCGAAGGCGTGTCCCTTGAC |
| **18** | Nup62-Flag-EPEA fwd | ACTTAAGCTTGGGCGATCGCAATGGCAAGCGGGTTTAATTTTGG |
| **19** | Nup62-Flag-EPEA rev | CTCTAGACTCGAGTTATGCTTCAGGTTCCTTATCGTCGTCATCCTTGTAGTCTG |
| **20** | SgfI-cJun fwd | CTGGCAGGCGATCGCAATGACTGCAAAGATGGAAACGACC |
| **21** | SgsI-cJun rev | TCTGGCGCGCCAAATGTTTGCAACTGCTGCGTTAGC |
| **22** | SgfI-nEPEA fwd | TACGCTGCGATCGCAATGGGCCAGCTGGTGGAGA |
| **23** | SgsI-nEPEA rev | CTGGCGCGCCAGAGCTCACAGTAACCTGGGTGCC |

**Supplementary References.**

1. Woo, C. M. *et al.* Development of IsoTaG, a Chemical Glycoproteomics Technique for Profiling Intact N- and O-Glycopeptides from Whole Cell Proteomes. *J Proteome Res* **16**, 1706-1718 (2017).
